# B cells dominate the heart immune infiltrate during chronic infectious cardiomyopathy

**DOI:** 10.64898/2026.11.09.765469

**Authors:** Aleksandra Dąbek, Amanda Fortes Francisco, Archie A. Khan, Shazia Ashraf, Shoumit Dey, Katrien Van Bocxlaer, Helen Ashwin, Kiera Parker, Nidhi S. Dey, Omer Ali Bayraktar, Sarah A. Teichmann, Martin C. Taylor, E. Allison Green, Paul M. Kaye, John M. Kelly, Michael D. Lewis, Damián Pérez-Mazliah

## Abstract

Recent advances in single-cell and spatially resolved multi-omics have revolutionised our understanding of the cellular and molecular niches governing cardiac homeostasis and acute injury. However, the immune-stromal networks sustaining long-term, chronic heart pathology remain poorly defined, largely due to a paucity of physiologically relevant experimental models. Here, through a side-by-side multi-omic comparison of *Trypanosoma cruzi*-infected mouse models displaying contrasting degrees of chronic myocardial pathology and fibrosis, we define the long-term cellular landscape of the inflamed fibrotic heart. We reveal that class-switched, antibody-secreting plasma B cells represent the predominant immune subset infiltrating the fibrotic myocardium. CellChat intercellular communication modelling identifies plasma cell-fibroblast crosstalk as the strongest immune-stromal interaction in the fibrosis-prone heart, driven by predicted matrix anchoring (CD44-collagen/laminin) and molecular survival axes (App-Cd74, C3-Cr2). Supporting these computational predictions, immunofluorescence and spatial profiling confirm close proximity between plasma B cells and fibroblasts, alongside a striking accumulation of B cells embedded within extracellular matrix-rich cardiac scars. This localised B cell response maintains prominent pools of parasite-reactive and heart-autoreactive antibody-secreting cells despite a lack of detectable parasite signal in their immediate micro-vicinity. Together, these findings identify a matrix-associated plasma B cell-fibroblast niche as a central driver of chronic cardiac remodelling, revealing how persistent tissue scarring can become decoupled from direct local pathogen presence.

## Introduction

Niches governing the healthy heart at steady state and during acute cardiac injury are slowly being unravelled^1–12^. Experimental mouse models of acute myocardial infarction and acute viral myocarditis have defined the transient immune cascades invading the damaged myocardium^5–7^, while contemporary human spatial atlases have begun translating these immune responses to clinical settings^5,8,10,13,14^. In contrast, the cellular and molecular mechanisms driving the progression and severity of chronic cardiac fibrosis and the immune-cell subsets contributing to chronic unresolving cardiomyopathy remain underexplored, largely due to a paucity of clinically relevant, long-term experimental animal models^15^. Addressing this gap is critical for uncovering novel immunopathological mechanisms and identifying targeted therapeutic interventions to halt progressive low-grade heart tissue scarring.

*Trypanosoma cruzi* infection, the cause of Chagas heart disease, represents the most common cause of chronic infectious cardiomyopathy globally and the highest-impact parasitic disease in the Americas^16–19^. Affecting an estimated 6–8 million people worldwide, with an additional 70 million at risk of transmission and over 12,000 annual deaths, Chagas disease has expanded from a historically neglected Latin American infection into a global health concern through international migration^18,20^. The initial acute phase of the disease is characterised by a high parasitaemia that is then controlled by the adaptive immune response, and those infected transition to the asymptomatic chronic stage, that is life-long in the majority of cases^21–23^. However, approximately one-third of infected individuals progress to chronic Chagas heart disease, frequently decades after the initial infection^24^. This chronic symptomatic phase is marked by myocardial inflammation, extensive interstitial fibrosis, and complex ventricular dysfunction^24–26^. No licenced vaccines are available, and current medicines are limited and often lead to considerable levels of adverse side-effects^24,27^. Crucially, while both parasite-specific protective immunity and cardiac-specific autoreactive immune responses have been described, how these dual immunological mechanisms interact to shape long-term tissue pathology and clinical outcome remains unresolved^24,28^. We have previously described different combinations of *T. cruzi* and mouse strains that model non-lethal chronic/persistent infections characterized by differing degrees of heart inflammation and fibrosis, which share a variety of features with human Chagas cardiomyopathy^29–31^. Thus, *T. cruzi* infection provides tractable experimental systems for deep phenotyping and molecular characterisation of chronic cardiomyopathy pathogenesis.

Here, to dissect the immune-stromal networks driving chronic cardiac inflammation and fibrotic remodelling, we utilised these *T.cruzi* infection mouse models^30^. Through integrated spatial transcriptomics, single-cell RNA sequencing, multiplexed imaging, and functional antibody-secretion assays, we map the long-term cellular architecture of the inflamed heart. Our findings uncover an unexpected cellular hierarchy, establishing a dominant role for matrix-anchored plasma B cells engaged in direct molecular crosstalk with cardiac fibroblasts, alongside cytotoxic NKG7+ T and NKT cells as essential orchestrators of chronic cardiac inflammation and fibrosis.

## Results

### Establishing a model for transcriptomics of immune responses in the heart during chronic cardiomyopathy

To investigate differing degrees of chronic cardiac pathology, we selected two *T. cruzi* infection models: C3H/HeN mice infected with *T. cruzi* strain JR (discrete typing unit (DTU) I) and BALB/c mice infected with *T. cruzi* strain CL Brener (DTU VI)^30^ (**Fig. 1A**). Parasites were engineered to constitutively express a red-shifted luciferase, enabling sensitive *in vivo* and *ex vivo* quantification of whole-body and tissue-specific parasite loads^30^. Both models are characterised by an acute phase with peak whole-body parasite burdens at approximately 2-3 weeks post infection (p.i.), which resolves over 4–6 weeks to establish chronic infection with sustainably low parasite loads^29,30^. Despite these similarities in acute-phase kinetics, C3H/HeN mice infected with *T. cruzi* JR develop significantly more severe cardiac fibrosis, with higher collagen deposition distributed more extensively between the muscle fibers compared to infected BALB/c mice^30^.

**Figure 1.**
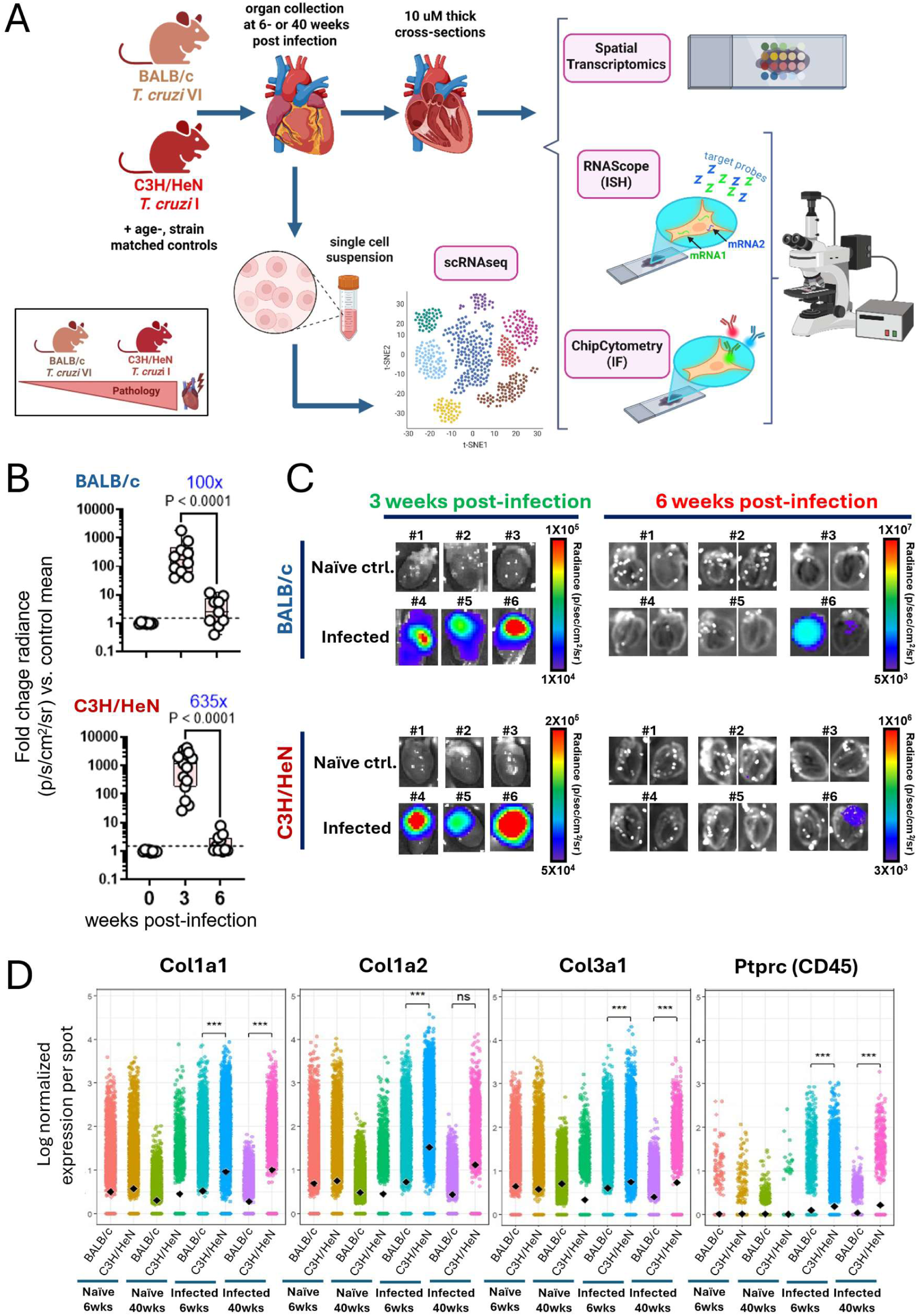
Experimental workflow, parasite burden, and spatial profiling of cardiac fibrosis and inflammation during chronic *T. cruzi* infection. **A)** Schematic overview of the experimental workflow. **B)** Fold-change in cardiac parasite radiance (p/s/cm²/sr) in BALB/c and C3H/HeN mice at 3 and 6 weeks p.i. relative to strain-, age-matched controls; statistical significance was assessed using an unpaired t-test. **C)** Representative bioluminescence images showing *T. cruzi* signal intensity in BALB/c and C3H/HeN hearts at 3 weeks (whole-heart imaging) and 6 weeks (imaged as two separate heart halves), alongside age- and strain-matched naïve controls. **D)** Quantification of normalised spot-level expression of *Col1a1/Col1a2*, *Col3a1*, and *Ptprc* at 6 weeks (n = 4 per condition) and 40 weeks (n = 2 per condition) p.i. in naïve and infected BALB/c and C3H/HeN mice. Data are shown as individual spatial transcriptomic spots with mean values (◆); statistical significance was determined using pairwise proportion tests (ns = not significant; *** = p < 0.0001).

To investigate the *in situ* immune response in the hearts from these two models, we performed integrated imaging and transcriptional analyses on C3H/HeN and BALB/c mice comparing the acute-to-chronic transition/early chronic (6 weeks p.i.) and late chronic (40 weeks p.i.) *T. cruzi* infection (**Fig. 1A**). By 6 weeks p.i., the cardiac parasite burden had declined substantially in both models. C3H/HeN mice showed a 635-fold reduction while BALB/c showed a 100-fold reduction compared to the 3 weeks p.i. time point, falling below the bioluminescence detection limit in approximately 50% of the C3H/HeN and BALB/c infected mice studied (**Fig. 1B-C, Suppl. Fig. 1, Suppl. Fig. 2A**).

Visium spatial transcriptomics (10x Genomics) confirmed that the expression of collagen mRNA (*Col1a1*, *Col1a2*, and *Col3a1*) and the pan-leukocyte marker CD45 (*Ptprc*) were significantly upregulated in the hearts from *T. cruzi*-infected C3H/HeN mice compared to infected BALB/c mice, both at 6 weeks and 40 weeks p.i. (**Fig. 1D** and **Suppl. Fig. 2**). Nonetheless, by 6 weeks p.i., both strains showed conspicuous collagen and *Ptprc* expression in the ventricle and atrium. While *T. cruzi*-infected C3H/HeN mice exhibited patches of very strong *Col1a1* and *Col1a2* expression in both the ventricle and atrium, the pattern of collagen expression in hearts from *T. cruzi*-infected BALB/c mice was spatially homogeneous. In agreement with our previous histopathology data showing no correlation between inflammation and fibrosis^30,32^, the expression of collagen genes was not directly associated with the expression of the *Ptprc* pan-leukocyte marker (**Supp. Fig. 2B**).

**Figure 2.**
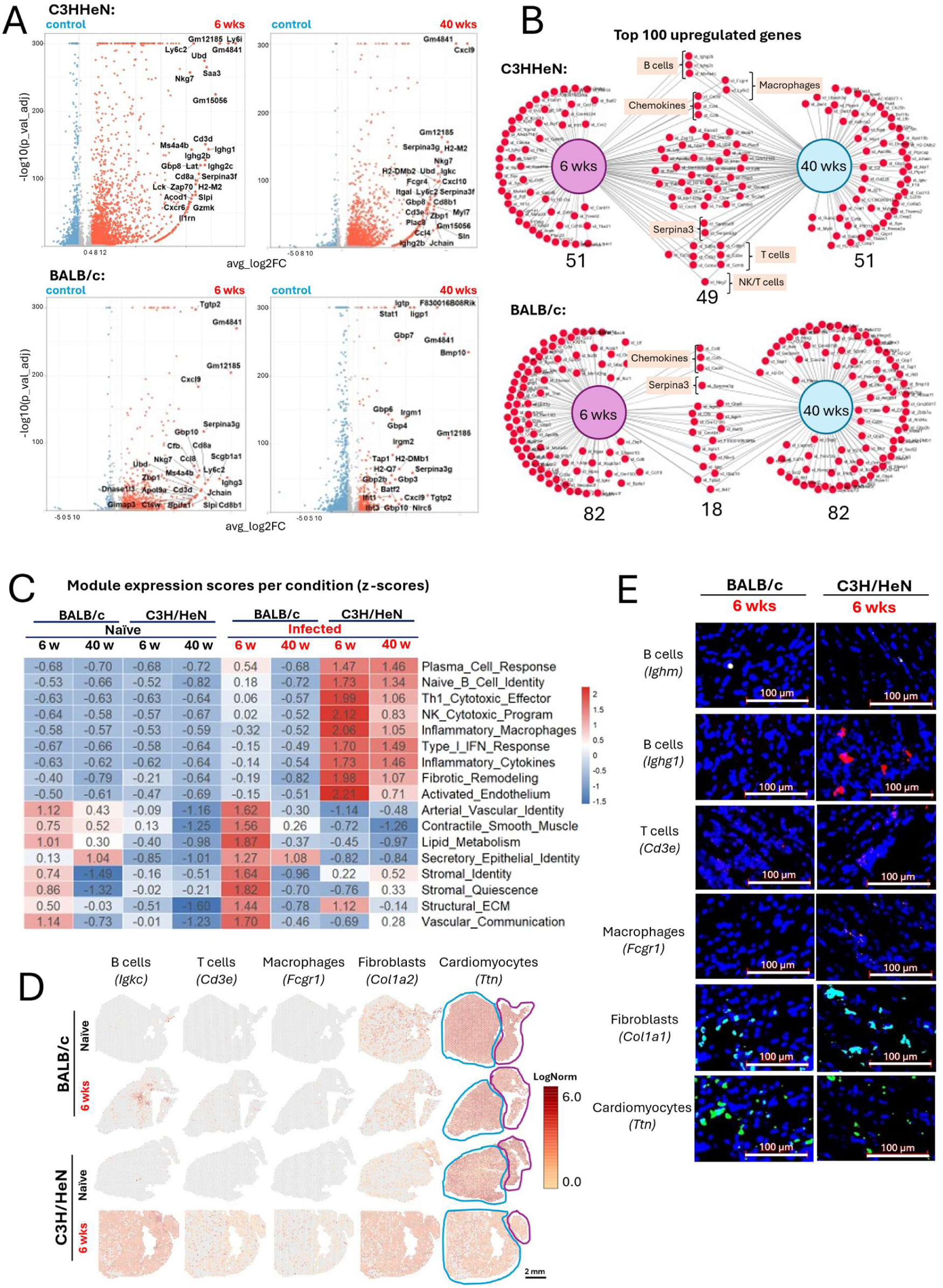
Transcriptional signatures of cardiac immune activation during early and late chronic infection. **A)** Volcano plots showing differential gene expression in spatial transcriptomic data between control and infected hearts at 6 weeks (n = 4 per condition, pooled from two independent experiments) and 40 weeks (n = 2 per condition) p.i. The top 25 upregulated genes (ranked by log₂ fold-change) are labelled. **B)** Venn diagrams illustrating the overlap between the top 100 upregulated genes identified at 6 and 40 weeks p.i. **C)** Heatmap displaying module expression z-scores derived from spatial transcriptomic data across conditions at both time points for selected biological processes. **D)** Spatial transcriptomic maps showing the expression and localisation of selected immune-related and structural genes at 6 weeks p.i. compared with age- and strain-matched controls. The blue dotted lines represent the ventricle boundaries, and the pink dotted lines represent the ventricle boundaries. **E)** Representative RNAscope images demonstrating mRNA expression of selected genes in cardiac tissue at 6 weeks p.i.

To determine whether local parasite burden drives spatial immune-stromal niche formation, we aligned spatial transcriptomics datasets against strain-matched *T. cruzi* reference genomes (Dm28c for TcI-infected C3H/HeN hearts; TCC for TcVI-infected BALB/c hearts, Materials and Methods, Spatial Transcriptomics). This analysis revealed no evidence of localised parasite transcription across tissue sections; detected parasite gene counts were extremely low (1 count per spot) and present equally in infected and uninfected control hearts, indicating non-specific background alignment. Thus, local parasite transcript accumulation is either very low or absent within the spatial niches occupied by either fibroblasts or infiltrating immune cells.

### Diversity of the heart immune infiltrate during chronic Chagas cardiomyopathy

We further used our spatial transcriptomics datasets to catalogue in greater detail the cardiac immune response during early and late chronic *T. cruzi* infection in C3H/HeN and BALB/c mice. Although both strains showed a striking immune infiltrate signature in the heart, C3H/HeN mice exhibited a broader and more sustained cardiac transcriptional response than BALB/c mice. At 6 weeks p.i., pseudo-bulk RNAseq differential gene expression analysis identified prominent upregulated B-cell-associated transcripts, including *Igkc*, *Ighg2b*, *Ighg2a*, *Ighg3*, *Ighg1* and *Jchain*, together with macrophage-associated genes (*Ly6c2*, *Fcgr4*, and *Itgal*) and T- and NK-cell-associated genes (*Nkg7*, *Cd3d*, *Cd3e*, *Cd3g*, *Cd8a*, and *Cxcr6*) in both mouse strains (**Fig. 2A-B**). While this striking transcriptional response remained evident at later time points in the heart from C3H/HeN mice, it was markedly attenuated in BALB/c hearts at 40 weeks (**Fig. 2B**). In contrast, a marked induction of inflammatory chemokines (*Ccl4*, *Ccl5*, *Ccl8*, *Cxcl9*, *Cxcl10*) and immune-activation *Serpina3g* genes remained evident at 6 and 40 weeks p.i. in both mouse strains (**Fig. 2B**).

Thus, comparison of the top upregulated genes at 6 and 40 weeks demonstrated a persistent core transcriptional program in C3H/HeN hearts, encompassing B-cell, macrophage, T-cell/NK-cell, chemokine, and *Serpina3*-associated genes. BALB/c mice showed substantially less overlap between the two time points, consistent with resolution or spatial restriction of the inflammatory response during late chronic infection (**Fig. 2B**). This sustained unresolved immune signature in the C3H/HeN model may be driven by higher frequencies of heart-specific parasitism as compared to BALB/c mice at these late time points p.i. ^30^.

Gene-module analysis further distinguished infected C3H/HeN hearts by enrichment of plasma B-cells, Th1/cytotoxic T-cells, NK-like T-cells, inflammatory macrophages, and type I interferon-response (IFN) programmes, both at 6 and 40 weeks p.i. (**Fig. 2C**). Modules related to endothelial activation, fibrosis, extracellular-matrix organisation, and vascular signalling were also altered, linking the inflammatory cardiac environment in C3H/HeN mice with transcriptional features of tissue remodelling (**Fig. 2C**). While the plasma B cell module was noticeably enriched at 6 weeks p.i. in BALB/c mice (albeit to a lower extent as compared to C3H/HeN mice), it resolved to baseline levels by 40 weeks p.i. (**Fig. 2C**). In contrast to the immune-dominant transcriptional profile of infected C3H/HeN hearts, BALB/c hearts at 6 weeks p.i. showed relative enrichment of vascular identity, smooth-muscles, metabolic-homeostasis, stromal-regulatory, transcriptional-stability, extracellular-matrix structural-identity, and vascular-signalling modules. This pattern is in line with preservation of tissue-resident vascular and stromal programs and a comparatively restrained inflammatory remodelling response as compared to C3H/HeN heart (**Fig. 2C**).

The module-based analysis was supported by Gene Ontology enrichment of differentially expressed genes at 6 weeks p.i. (**Suppl. Fig 3**). In C3H/HeN hearts, enrichment of adaptive and innate immune-response, leukocyte-activation, T-cell-activation, and cytokine-production terms paralleled the increased immune-cell, interferon-associated, endothelial-activation, and fibrosis-related module scores (**Fig. 2C** and **Suppl. Fig 3**). Strikingly, C3H/HeN mice showed GO enrichment of B cell activation, humoral immune responses and immunoglobulin production (**Suppl. Fig 3B**). Conversely, BALB/c hearts showed enrichment of mitochondrial respiratory-chain assembly, NADH-dehydrogenase-complex assembly, fatty-acid metabolic processes, and VEGF signalling, in line with their relative enrichment for metabolic-homeostasis and vascular-signalling modules (**Fig. 2C** and **Suppl. Fig 3**).

Spatial mapping of representative lineage markers revealed that the B-cell-associated transcript *Igkc* was the most prominent immune signal within hearts from infected mice (**Fig. 2D**). In BALB/c mice at 6 weeks p.i., *Igkc* expression was localised to discrete focal regions mainly within the ventricle, whereas C3H/HeN hearts showed a much wider distribution throughout the ventricle and atrium (**Fig. 2D**). T-cell (*Cd3e*) and macrophage (*Fcgr1*) signals were also detected but were less spatially prominent than the B-cell-associated signal (**Fig. 2D**). Interestingly, the increased immune signature in hearts from *T. cruzi*-infected C3H/HeN was associated with a relative decrease in the expression of the cardiomyocyte-associated *Ttn* gene (**Fig. 2D**).

RNAscope imaging at 6 weeks p.i. confirmed the presence of B cells, T cells and macrophage-associated transcripts within cardiac tissue from infected mice (**Fig. 2E**), as well as the pattern of expression of fibroblasts-associated *Col1a1* and cardiomyocytes-associated *Ttn* genes. Notably, immunoglobulin-associated B-cell signals, particularly *Ighg1*, were more prominent in C3H/HeN than in BALB/c hearts, corroborating the spatial transcriptomic evidence for a B cell–enriched infiltrate in the more severe fibrotic-prone strain (**Fig. 2E**).

Together, these analyses indicate that C3H/HeN hearts primarily undergo an inflammatory transcriptional response, with a striking B cell response that is spatially widespread in both the atrium and ventricle *in situ* in the heart. In contrast, BALB/c hearts retain a comparatively vascular- and metabolism-associated tissue program during chronic *T. cruzi* infection, with highly localised areas of B cell enrichment during early chronic infection that resolve at later time points.

To validate the immune-cell distributions identified by spatial transcriptomics at the protein level, we performed ChipCytometry CellScape multiplex immunofluorescence on heart sections collected at 6 weeks p.i., a time point at which prominent immune responses were readily detected in both *T. cruzi*–infected C3H/HeN and BALB/c mice. For this, we used an 18-plex panel (**Table 2**) which enabled single-cell spatial quantification of leukocytes and major lymphoid and myeloid populations, including CD4+ and CD8+ T cells, regulatory T cells (Tregs), B cells, dendritic cells (DCs), monocytes/macrophages, neutrophils, and fibroblasts (**Tables 2 and 3**).

Consistent with the spatial-transcriptomic findings, C3H/HeN hearts showed a pronounced inflammatory infiltrate relative to strain-matched naïve controls (**Fig. 3**). This was characterised by significant increases in IgG+ B cells, CD8+ T cells, CD4+ T cells, and F4/80+ monocytes/macrophages. IgG^+^ B cells were the most abundant infiltrating population in the heart from *T. cruzi*-infected C3H/HeN mice, with a median density of approximately 900 cells/mm², followed by CD8+ T cells (∼700 cells/mm²), CD4+ T cells (∼300 cells/mm²), and F4/80+ macrophages (∼150 cells/mm²). Strikingly, these heart-resident B cells showed no expression of CD19 or CD138, classic mouse B cell and plasma cell markers, respectively. Representative multiplex images confirmed the accumulation of these populations within infected C3H/HeN myocardial sections (**Fig. 3 A-D**). In contrast, BALB/c hearts displayed a substantially attenuated cellular infiltrate at the same time point (**Fig. 3**). Among the populations quantified, only CD8+ T cells were significantly increased relative to naïve BALB/c controls, and their density was markedly lower than that observed in infected C3H/HeN hearts (approximately 20 cells/mm²). Although B cells, CD4+ T cells, and macrophages were detectable in infected BALB/c hearts, their increases did not reach statistical significance (**Fig. 3 A-D**).

**Figure 3.**
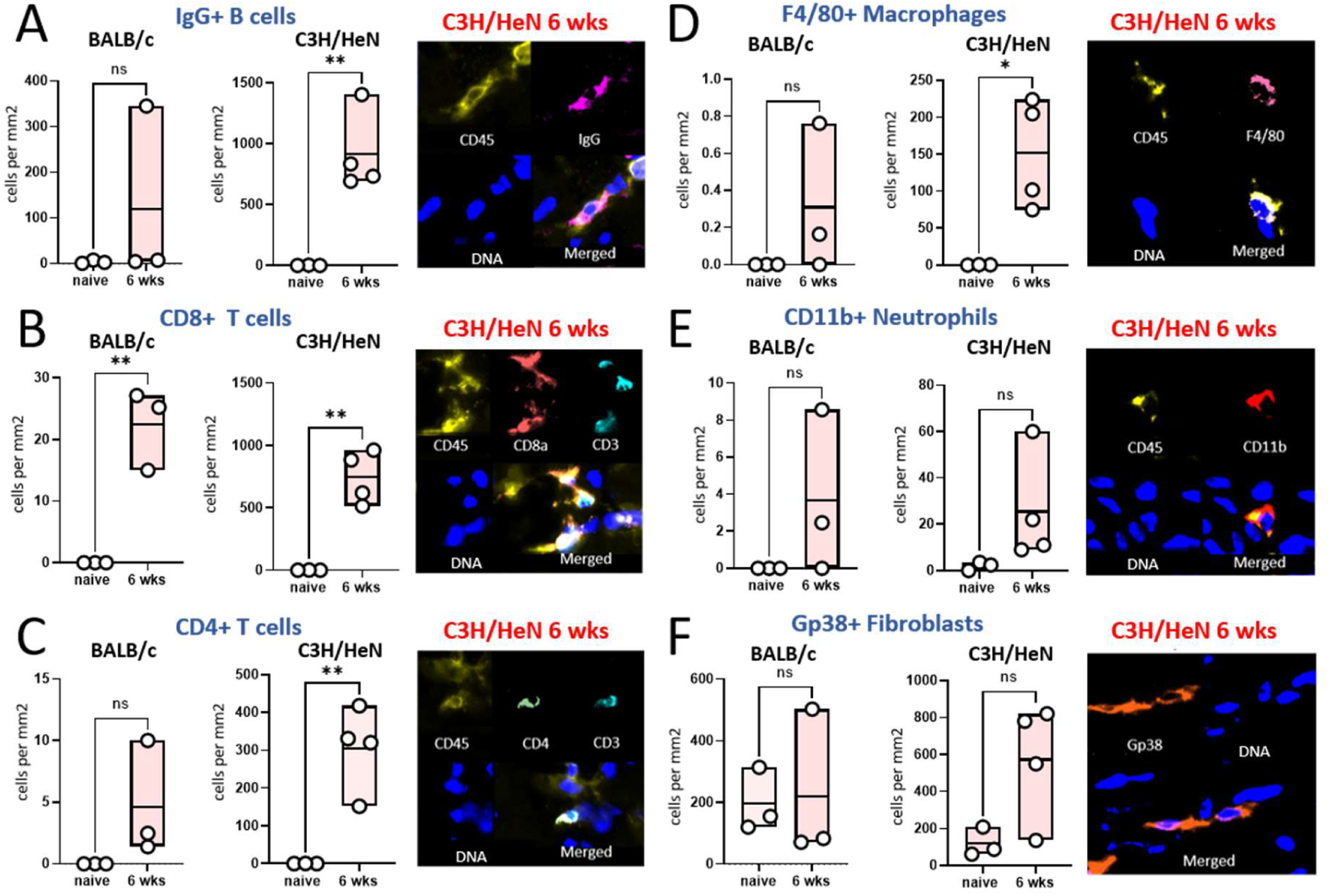
ChipCytometry analysis of immune and stromal cell populations in the heart. Quantification of **A)** IgG⁺ B cells, **B)** CD8⁺ T cells, **C)** CD4⁺ T cells, **D)** F4/80⁺ macrophages, **E)** CD11b⁺ neutrophils, and **F)** Gp38⁺ fibroblasts per mm² in 10 uM heart cross-sections from BALB/c and C3H/HeN mice, together with representative immunofluorescence (IF) images of each cell population. Statistical comparisons between BALB/c 6-week and C3H/HeN 6-week groups were performed using two-tailed unpaired Student’s *t*-tests. Individual data points are overlaid on box-and-whisker plots; the centre line indicates mean value.

Neutrophil, DCs and fibroblast densities did not differ significantly between infected and naïve animals in either strain (**Fig. 3 E-F** and data not shown). In addition, FoxP3^+^ Tregs were not detected in any conditions. Together, these protein-level data support the spatial transcriptomics evidence of a stronger and more diverse cardiac immune response dominated by B cells in C3H/HeN mice, whereas BALB/c mice exhibited a comparatively limited infiltrate dominated by CD8+ T cells.

To define the transcriptional states of infiltrating immune cells at single-cell resolution, we performed scRNA-seq on cardiac tissue from naïve and *T. cruzi*–infected BALB/c and C3H/HeN mice at 6 weeks p.i. Following quality control and integration, 129, 326 cardiac cells were resolved into major stromal, vascular, myocardial, and immune populations, including B cells, T cells, macrophages, dendritic cells, neutrophils, and proliferating lymphoid and myeloid populations (**Fig. 4 A–C**). Immune-cell representation was increased in infected hearts, particularly in C3H/HeN mice, aligning with the stronger inflammatory infiltrate detected by spatial transcriptomics and CellScape ChipCytometry.

**Figure 4.**
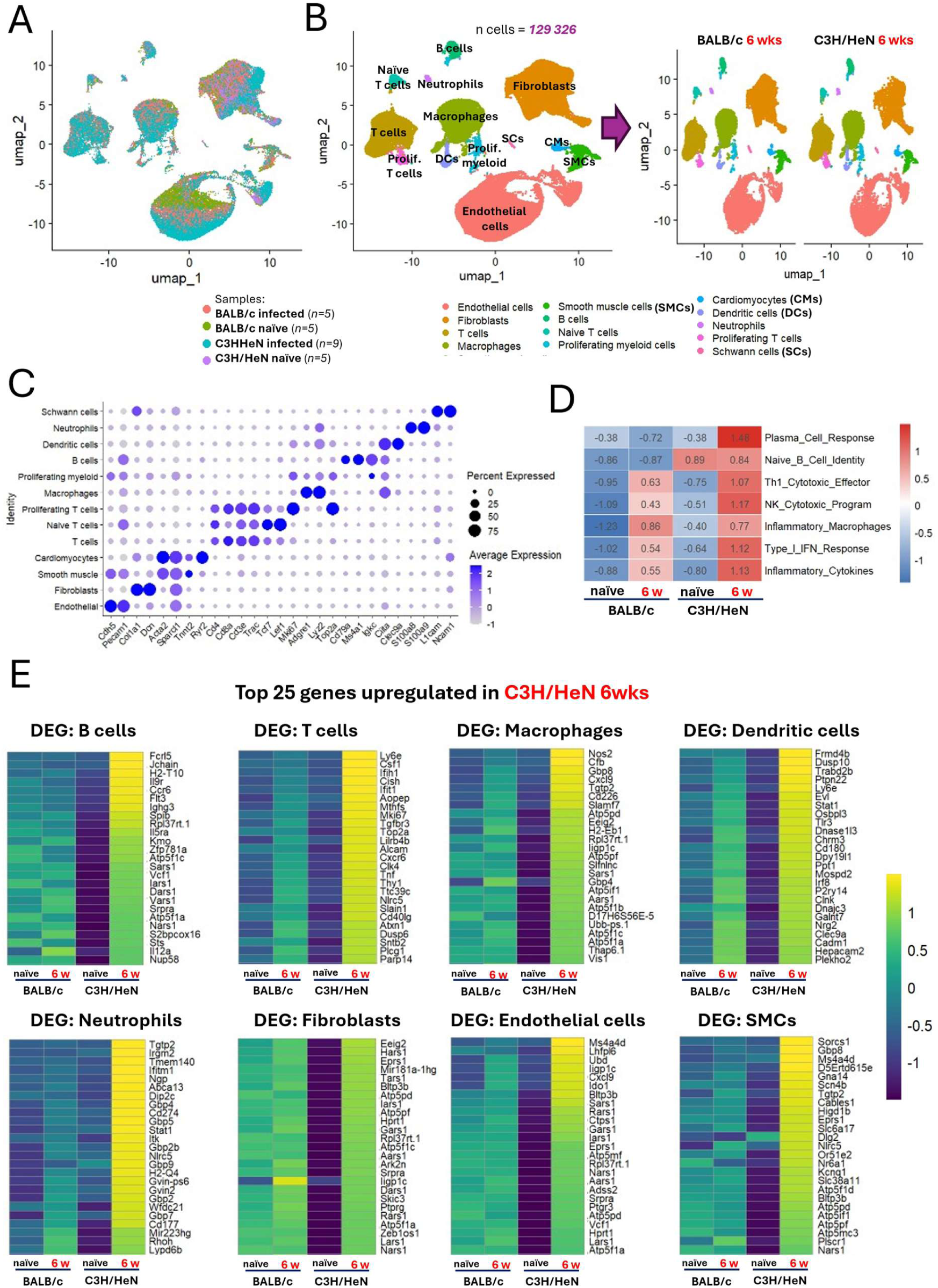
Single-cell profiling of cardiac immune and stromal populations during early chronic infection. **A)** UMAP embedding of cardiac scRNA-seq data showing sample distribution grouped by condition (BALB/c and C3H/HeN mice at 6 weeks p.i., n=5 and 9 respectively, together with age- and strain-matched controls, n=5 per strain). **B)** Annotated UMAP identifying major immune and stromal cell populations isolated from cardiac tissue. **C)** Heatmap displaying the expression of canonical marker genes used to define each cell-type cluster. **D)** Heatmap displaying module expression z-scores derived from scRNA-seq data across conditions at both time points for selected biological processes. **E)** Heatmaps showing the top 25 highest z-scores in selected cell types for C3H/HeN at 6 weeks p.i. compared with other conditions. Genes were selected from the top 50 upregulated genes in C3H/HeN 6 weeks relative to C3H/HeN naïve.

Module analysis showed that plasma B-cell signatures were among the most strongly enriched immune programs at 6 weeks p.i., with the greatest enrichment observed in C3H/HeN hearts (**Fig. 4D**). This transcriptional pattern was notable despite the relatively modest representation of B-lineage cells in the scRNA-seq dataset compared with the extensive B-cell infiltration indicated by 10X Visium, RNAScope and CellScape chipcytometry (**Fig. 4 A–C**). This lower B cell abundance in the scRNA-seq dataset likely reflects known technical challenges in isolating delicate antibody-secreting plasma cells during tissue dissociation^8,9,33–35^. Moreover, hearts from *T. cruzi*-infected C3H/HeN mice exhibited significant enrichment in various inflammatory gene expression modules, such as Th1 cytotoxic T cells, NKT cells, inflammatory cytokines, and a marked Type I IFN response, when compared to hearts from *T. cruzi*-infected BALB/c mice (**Fig. 4D**). Interestingly, the inflammatory macrophage module was enriched in hearts from both *T. cruzi*-infected C3H/HeN and BALB/c mice (**Fig. 4D**). In agreement with spatial data sets, DCs constituted a minor fraction of the overall cardiac cell population in both strains (**Fig. 4B**). However, they were more prominently represented in the hearts from *T. cruzi*-infected C3H/HeN mice within the scRNA-seq data set. Likewise, neutrophils were found to be a scarce population in the hearts from both BALB/c and C3H/HeN mice 6 weeks p.i. (**Fig. 4B**).

**Fig. 4E** details the top 25 upregulated transcriptional profiles in cardiac immune cell populations from infected C3H/HeN mice (at 6 weeks p.i.) compared with naive controls and infected BALB/c mice. Within the B-cell compartment, *Fcrl5* and *Jchain* were the two most highly upregulated genes in C3H/HeN mice. *Jchain* expression is among the best markers consistent with immunoglobulin-secreting plasma cells/plasmablasts^36^, whereas *Fcrl5* identifies recently-activated antigen-experienced B-cells^37^. This was accompanied by upregulation of heavy and light chain immunoglobulin genes (*Ighg3*, *Igkc*, *Iglc2*), together with activation, class switching, homing and cytokine receptor genes (*H2-T10*, *Il9r*, *Ccr6*, *Flt3*, *Rpl37rt*, *Il5ra*, and *Zfp781a*). This reinforces a distinct humoral signature with accumulation of recently-activated B cells and their differentiation into antibody-secreting cells in hearts from *T. cruzi*-infected C3H/HeN mice.

T cells from *T. cruzi*-infected C3H/HeN hearts exhibited a strong proliferative, interferon-responsive, and cytotoxic/activation profile. Key upregulated genes included cell cycle/proliferation markers (*Mki67*, *Top2a*), type I IFN genes *(ISGs*) (*Ifit1*, *Ifih1*, *Ly6e*), and key activation/cytotoxic molecules (*Cd8a*, *Cxcr6*, *Tnf*, *Cd40lg*, *Nlrc5*) (**Fig. 4E**). Interestingly, and despite their relatively low abundance, DCs from *T. cruzi*-infected C3H/HeN hearts show a distinct antigen cross-presentation, activation, and ISG signature. Top marker features include *Stat1*, *Tlr3*, *Irf8*, *Clec9a*, *Xcr1*, *Cd180*, and *Ly6e* (**Fig. 4E**).

Macrophages were abundant in hearts from both *T. cruzi*-infected C3H/HeN and BALB/c mice. However, in comparison, they displayed a dominant pro-inflammatory/M1-skewed activation signature in C3H/HeN hearts. Key features include elevated expression of *Nos2*, *Cxcl9*, complement factor B (*Cfb*), interferon-inducible GTPases (*Gbp8*, *Tgtp2*, *Gbp4*, *ligp1c*), and antigen presentation/co-stimulation genes (*H2-Eb1*, *Slamf7*, *Cd226*) (**Fig. 4E**). Side-by-side comparison of cardiac macrophages at 6 weeks p.i. in C3H/HeN vs BALB/c mice revealed stark transcriptional differences (**Suppl. Fig 4**). Cardiac macrophages from *T. cruzi*-infected C3H/HeN mice were heavily skewed toward pro-inflammatory, M1-associated, type I/II interferon-stimulated, and complement-related genes, including *Stat1*, *Nos2*, *Cxcl9*, *Ccl8*, *Tnf*, *Ifit3*, *Irf7*, *Rsad2*, *Cd274 (PD-L1)*, *C1qa*, *C1qb*, *Psmb8*, and *Fcer1g* (**Suppl. Fig 4A**). On the other hand, cardiac macrophages from *T. cruzi*-infected BALB/c mice retained higher relative expression of anti-inflammatory, tissue repair, and homeostatic/regulatory markers, including *Cd163*, *Arg2*, *Socs3*, *Hpgd*, *Pla2g7*, *Apoe*, and *S100a9* (**Suppl. Fig 4A**). In addition, gene ontology **(GO)** analysis showed strong enrichment of cytotoxic, inflammatory, and interferon-driven programmes, including cellular response to type II interferon (IFN-γ), cellular response to LPS / bacterial molecules, cell killing, leukocyte/lymphocyte proliferation, and NK cell-mediated immunity pathways in cardiac macrophages from *T. cruzi*-infected C3H/HeN mice (**Suppl. Fig 4B**). Conversely, cardiac macrophages from *T. cruzi*-infected BALB/c mice showed enrichment in tissue maintenance and vascular communication pathways, including endothelial/epithelial cell migration, tRNA aminoacylation, and mitochondrial ATP synthesis / oxidative phosphorylation (**Suppl. Fig 4B**). To better resolve the functional states of infiltrating and tissue-resident macrophages in the heart from both mouse strains, we re-clustered the cardiac macrophage compartment and applied UCell^38^ module scoring based on canonical functional gene sets. This analysis identified eight distinct macrophage sub-clusters spanning six major operational states: Resident 1/ Resident 2 (*Nr3c2*⁺/*Cpne8*⁺/*Cd163*⁺), Regenerative (*Tgfb3*⁺), Inflammatory (*Trem1*⁺/*Plac8*⁺/*Sell*⁺), Recruited (*Ccr2*⁺), Cycling/Proliferative (*Kif18b*⁺/*Ccna2*⁺), and Complement-expressing macrophages, which resolved into Complement-Resident (*Cfp*⁺/*Pltp*⁺) and Complement-Inflammatory sub-states (*C3*⁺/*Cxcl9*⁺/*Cxcl10*⁺) (**Suppl. Fig 4 C-F**). Comparing macrophage state proportions at 6 weeks p.i. revealed striking strain-specific divergences (**Suppl. Fig 4 C**). In hearts from infected BALB/c (BALB/c 6wks), the macrophage response was marked by a heavy enrichment of *Ccr2*⁺recruited macrophages, alongside the retention of Regenerative and Resident subsets (**Suppl. Fig 4 C** and **E**). In stark contrast, fibrosis-prone C3H/HeN hearts (C3H/HeN 6wks) underwent a massive shift toward Complement-expressing (both resident and inflammatory) as well as proliferative macrophages (**Suppl. Fig 4 C** and **E**). Notably, complement-expressing macrophages represented the primary macrophage source of the lymphocyte chemoattractant *Cxcl9* and *Cxcl10*.

Together, these findings show that chronic infection in C3H/HeN mice drives a profound expansion of complement- and chemoattractant-producing *Nos2*⁺ M1-like pro-inflammatory cardiac macrophages that sustain cytotoxic lymphocyte recruitment, whereas BALB/c hearts maintain a balanced myeloid landscape dominated by *Ccr2*⁺ recruited and tissue-resident cardiac macrophages that likely help preserve myocardial structural integrity.

Interestingly, both spatial transcriptomics and scRNA-seq revealed that hearts from *T. cruzi*-infected C3H/HeN mice exhibited a striking Type I IFN gene signature at 6 weeks p.i. (**Fig. 2C** and **4D**, **Suppl. Fig. 5A-B**), which remained sustained at later 40-week p.i. time points (**Fig. 2C**, **Suppl. Fig. 5B**). Albeit to a lower magnitude, this Type I IFN signature was also detected and maintained during late chronic stages in hearts from *T. cruzi-*infected BALB/c mice (**Suppl. Fig. 5B**). Cardiac spatial distribution of the signature showed higher compartmentalisation and restriction to the atria in *T. cruzi*-infected BALB/c mice, as opposed to an even distribution among atria and ventricles in *T. cruzi*-infected C3H/HeN mice (**Suppl. Fig. 5E**).

Single-cell profiling at 6 weeks p.i. revealed that interferon-stimulated genes (ISGs) were prominent top marker genes across nearly every immune cell compartment in the hearts from *T. cruzi*-infected C3H/HeN mice. This included T cells (*Ifit1*, *Ifih1*, *Ly6e*), macrophages (*Stat1*, *Ifit3*, *Irf7*, *Rsad2*), DCs (*Stat1*, *Ly6e*, *Irf8*), and neutrophils (*Tgtp*2, *Ifitm1*, *Stat1*, *Zbp1*) (**Fig. 4E**). Neutrophils and DCs exhibited the highest relative per-cell expression density for core ISGs in both mouse strains (e.g., *Ifit3*, *Zbp1*, *Gbp5*, *Oasl1/2* in neutrophils; *Irf8* in DCs), followed by macrophages and, to a lesser extent, T cells (**Suppl. Fig. 5C-D**). Non-immune structural and vascular cells, most notably endothelial cells (ECs), Schwann cells (SCs), and fibroblasts, also participated in the Type I IFN programme, expressing elevated levels of ISGs such as *Ifit2*, *Gbp5*, and *Rsad1* (**Suppl. Fig. 5C-D**).

Thus, both *T. cruzi*-infected BALB/c and C3H/HeN mice exhibited a distinctive cardiac Type I IFN response. However, in C3H/HeN hearts, myeloid and granulocytic populations reached extreme peak z-scores (e.g., *Ifit3* and *Oasl2* exceeding values of +3.0 to +4.0), which could lead to a much higher local concentration of interferon signalling compared to cardiac BALB/c myeloid cells (**Suppl. Fig. 5D**).

### Stromal, Vascular, and Metabolic Homeostasis in the heart during *T. cruzi* infection

Our single-cell datasets allowed us to transcriptionally characterise endothelial cells (ECs), smooth muscle cells (SMCs) and fibroblasts in hearts from *T. cruzi*-infected BALB/c and C3H/HeN mice (**Fig. 4** and **Suppl. Fig. 6 and 7**). At the global tissue level, fibroblasts, ECs, and SMCs in hearts from infected BALB/c mice mount a strikingly coordinated, multi-lineage gene program dominated by two gene families: MHC and *Serpina3* (**Suppl. Fig. 6**). All three cell populations showed upregulation of H2/MHC genes, including classical MHC Class I (*H2-K2*, *H2-D1*), non-classical MHC Class I (*H2-Q5*, *H2-Q6*, *H2-T15*), and MHC Class II subunits (*H2-Ea*, *H2-Eb1*). There was also upregulation of Gbp genes (*Gbp2*, *Gbp3*, *Gbp4*, *Gbp5*, *Gbp6*) and other interferon-stimulated transcripts (*Tgtp1*, *Tgtp2*, *Tap1*, *Zbp1*, *Ifi47*), alongside co-induction of CXCR3-binding chemokines (*Cxcl9*, *Cxcl10*, *Cxcl11*). Additionally, all three cell populations showed prominent expression of the *Serpina3* family genes (*Serpina3g*, *Serpina3h*, *Serpina3h.1*, *Serpina3i*, *Serpina3n*), which are serine protease inhibitors that neutralise tissue-destructive enzymes such as cathepsins, elastases and granzymes^39^ (**Suppl. Fig. 6**). A seemingly coordinated gene programme was also detected in fibroblasts, ECs, and SMCs within hearts from infected C3H/HeN mice. However, in contrast with BALB/c mice, the programme in C3H/HeN mice seemed strongly biased towards a hyper-metabolic, ATP-driven bioenergetic signature, characterised by a striking enrichment of mitochondrial ATP synthase complex subunits, including *Atp5f1a*, *Atp5f1b*, *Atp5f1c*, *Atp5pd*, and *Atp5pf* (**Fig. 4E**).

To better resolve cardiac fibroblast heterogeneity during *T. cruzi* infection, we performed UCell-guided sub-clustering^38^, which resulted into five distinct subclusters (**Suppl. Fig. 7A**). These were then annotated based on established adult cardiac fibroblast single-cell atlases^1,8,9,40,41^. At baseline, hearts from naive BALB/c mice were defined by three resting homeostatic populations: Baseline Interstitial Fibroblasts (F-Base-Int, *Ptger4*⁺/*Jup*⁺/*Junb*⁺/*Ccn1*⁺/*Col5a3*⁺), representing the dominant resting interstitial pool of the main myocardial wall^1,40^; ECM Structural Fibroblasts (F-Base-Str, *Nkain3*⁺/*Piezo2*⁺/*Smad*6⁺/*Igfbp3*⁺/*Eya4*⁺/*Pdlim3*⁺/*Apoe*⁺), which functions as a mechanosensitive and matrix-organising niche expressing inhibitory TGF-β regulators (*Smad6*)^42^; and a specialized population of Baseline WntX Fibroblasts (F-WntX, *Wif1*⁺/*Erbb4*⁺/*Kcnma1*⁺/*Clu*⁺/*Rspo3*⁺/*Dsp*⁺), localised to perivalvular and boundary niches, F-WntX fibroblasts function as a constitutive Wnt-inhibitory checkpoint through high expression of secreted Wnt antagonists (*Wif1*, *Rspo3*) and matrix chaperones (*Clu*)^41,43^ (**Suppl. Fig. 7A-C**). Following *T. cruzi* infection, hearts from BALB/c mice showed a regulated stromal remodelling response characterised by a decrease in baseline interstitial and structural fibroblasts, accompanied by the selective expansion of Activated Fibroblasts (F-Activ, *Gfra1*⁺/*Dpep1*⁺/*Slit3*⁺/*Pde4b*⁺/*Sod2*⁺/*Thsd7a*⁺/*Prex2*⁺) (**Suppl. Fig. 7A, D**). These fibroblasts upregulated genes implicated in stromal activation, cell and ECM adhesion and cell motility (*Dpep1*, *Slit3*, *Thsd7a*, *Prex2*) in combination with genes implicated in antioxidant defence and intracellular cAMP regulation (*Sod2*, *Pde4b*). In comparison, hearts from fibrosis-prone naïve C3H/HeN mice exhibited a pre-existing baseline deficit in interstitial structural fibroblasts, together with a relative enrichment of a complement-producing Type I/II IFN-responsive fibroblast population (F-C3-IFN, *C3*⁺/*Gbp7*⁺/*Gbp9*⁺/*Gbp2b*⁺/*Spon1*⁺/*Stat1*⁺/*Irf1*⁺/*Mt1*⁺/*Mt2*⁺) (**Suppl. Fig. 7A, D**). Upon *T. cruzi* infection, this F-C3-IFN population heavily accumulated, becoming the dominant fibroblast population in hearts from infected C3H/HeN mice (**Suppl. Fig. 7A, D**). Contrary to infected BALB/c mice, the F-Activ population described above remained close to baseline.

Thus, fibroblasts, ECs, and SMCs appeared to coordinate a dual pathogen-reactive response while also helping preserve cardiac extracellular matrix integrity during active inflammation in hearts from BALB/c mice, consistent with increased vascular, stromal, contractile, and structural preservation identity gene modules (**Fig. 2C**). In contrast, these cell populations seemed to actively contribute to the increased fibrotic remodelling and activated endothelium modules in hearts from *T. cruzi*-infected C3H/HeN mice (**Fig. 2C**).

### Plasma (B) cells dominate immune-fibroblast crosstalk in the heart during chronic Chagas cardiomyopathy

After broad in depth analysis of immune responses *in situ* in the heart during *T. cruzi*-infection, accumulation of a striking plasma (B) cell signature was a particularly striking feature of the more cardiac fibrosis-susceptible C3H/HeN model (**Fig. 2-4**). SPOTlight deconvolution^44^ showed that B cells represented the single most abundant cell type in these hearts, accounting for ∼23% of the total average cell proportion (**Fig. 5A-B**). In contrast, the relative abundance of B cells in hearts from *T. cruzi*-infected BALB/c remained below SMCs and ECs (**Fig. 5A-B**).

**Figure 5.**
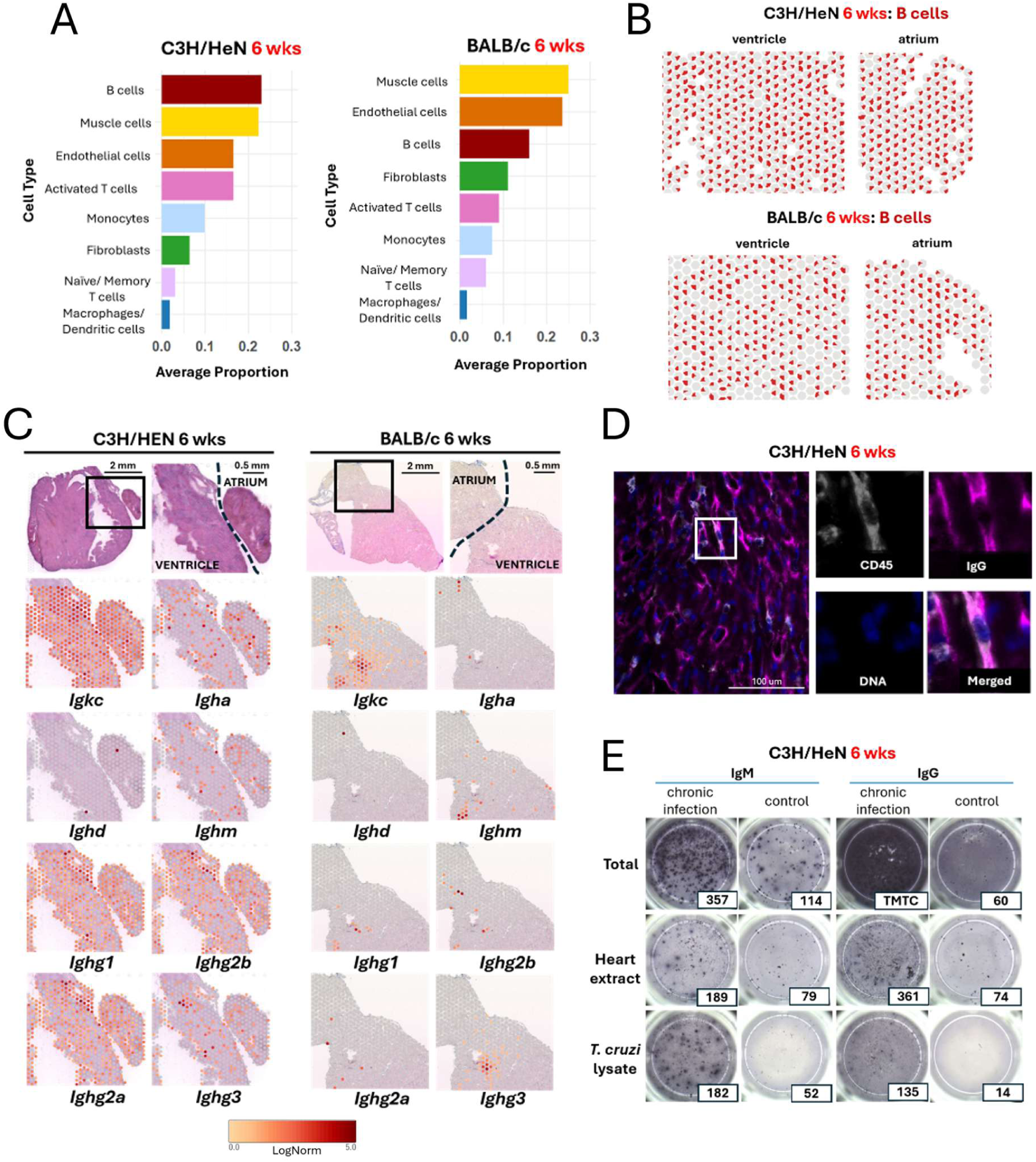
Spatial, protein, and functional profiling of cardiac B-cell responses during chronic infection. **A)** Bar plots showing predicted cell-type abundance derived from SPOTlight deconvolution of spatial transcriptomic data at 6 weeks p.i. in C3H/HeN and BALB/c mice. **B)** Representative spatial transcriptomic maps displaying the SPOTlight predicted abundance of B cells (red) across cardiac tissue sections from C3H/HeN and BALB/c mice. **C)** Spatial transcriptomic maps showing the distribution and expression intensity of immunoglobulin heavy-chain isotype genes across cardiac tissue sections during chronic infection in C3H/HeN and BALB/c mice. **D)** Immunofluorescence staining demonstrating IgG-expressing B-cell infiltration in chronically infected C3H/HeN mice. **E)** B-cell ELISpot assay quantifying antibody-secreting cells (ASCs) isolated from hearts of chronically infected C3H/HeN mice compared with naïve age- and strain-matched controls. Spots represent individual IgM- and IgG-secreting B cells (number of spots indicated).

Moreover, the spatial distribution of the cardiac B cell signature greatly differed between *T. cruzi*-infected C3H/HeN and BALB/c mice (**Fig. 5C**). Spatial maps showed that B cells were broadly distributed across both ventricular and atrial myocardial tissue in *T. cruzi*-infected C3H/HeN mice. In contrast, the B cell signal was confined to rare isolated focal patches mainly located in ventricular areas in the heart of *T. cruzi*-infected BALB/c mice (**Fig. 5C**). Spatial transcriptomic maps revealed intense expression of light-chain *Igkc* alongside extensive class-switched heavy-chain genes in hearts from *T. cruzi*-infected C3H/HeN mice. The infiltrating B-cell population in C3H/HeN hearts was heavily enriched for class-switched IgG isotypes, predominantly *Ighg1*, *Ighg2b*, *Ighg2a*, and, to a lesser extent, *Ighg3*. This was alongside *Ighm* and, unexpectedly, a distinct widespread *Igha* signal, albeit to a lesser extent as compared to IgG isotypes (**Fig. 5C**). In agreement, immunofluorescence analysis detected a striking accumulation of DAPI^+^CD45^+^IgG^+^ B cells in the heart from *T. cruzi*-infected C3H/HeN mice (**Fig. 5D** and **3A**). In contrast, *Ighg3* was the most prominent IgG isotype in hearts from *T. cruzi*-infected BALB/c mice (**Fig. 5C**).

Functional B-cell ELISPOT assays on single-cell cardiac suspensions verified that B cells infiltrating the hearts from *T. cruzi*-infected C3H/HeN mice were actively secreting antibodies (**Fig. 5E**). Our data confirmed robust IgG-producing responses, directed both against parasites and against self-cardiac antigens. Specifically, parasite-reactive antibody-secreting cells were markedly elevated, generating 182 IgM spots against crude *T. cruzi* antigen extract versus 52 in naive controls and 135 IgG spots versus 14 in naive controls (**Fig. 5E**). Crucially, this parasite-directed response was accompanied by an influx of heart-autoreactive antibody-secreting cells targeting cardiac antigens, yielding 189 IgM spots versus 79 in naive controls and 361 IgG spots versus 74 in naive controls (**Fig. 5E**). Thus, cardiac B cell responses were broadly distinct between *T. cruzi*-infected C3H/HeN and BALB/c mice, differing in magnitude, isotype composition, spatial distribution and functional activity.

Interestingly, dual immunofluorescence staining and confocal microscopy showed that Podoplanin⁺ cardiac fibroblasts^45^ reside in close physical proximity to infiltrating IgG⁺ B cells/plasma cells in hearts from *T. cruzi*-infected C3H/HeN mice (**Fig. 6A**). Wheat Germ Agglutinin (WGA) counterstaining demonstrated that infiltrating IgG⁺ B cells/plasma cells were frequently embedded directly within regions of heavy extracellular matrix (ECM) deposition and fibrotic scarring in hearts from *T. cruzi*-infected C3H/HeN mice (**Fig. 6B**). These findings motivated the use of CellChat computational predictions to interrogate how these plasma B cells could interact with ECM and fibroblasts. Indeed, CellChat analysis on our scRNA-seq datasets identified a marked enrichment in overall communication strength between B cells and fibroblasts specifically in hearts from *T. cruzi*-infected C3H/HeN mice (**Fig. 6C**). In agreement with our immunofluorescence results (**Fig. 6A-B**) CellChat analysis identified strong matrix-to-lymphocyte interactions in the fibroblast to B cell direction, dominated by fibroblast-derived collagens (*Col1a1*, *Col1a2*, *Col4a1*, *Col6a3*) laminins (*Lama2*, *Lama4*, *Lamb1*, *Lamb2*, *Lamc1*) and fibronectin (*Fn1*) engaging CD44 and Syndecan-4 (*Sdc4*) on B cells (**Fig. 6D-E**). Although hyaluronic acid is the canonical ligand for CD44, CD44 can directly interact with fibronectin and establish secondary interactions with collagen and laminin matrix components, providing a mechanical and structural tether that retains infiltrating B cells/plasma cells within the newly synthesised fibrotic matrix. This B-cell-to-ECM interaction was reinforced by THBS gene family members (*Thbs1* and *Thbs2*, encoding a group of secreted matricellular glycoproteins that bridge cell-to-ECM) interacting with CD47 and Syndecan-4, and tenascin-X (*Tnxb*, encoding a large extracellular matrix glycoprotein) further interacting with Syndecan-4 on B cells (**Fig. 6D-E**). Importantly, CellChat also identified well-characterised stromal-immune interactions in the fibroblast to B cell direction, including a strong fibroblast amyloid precursor protein (*App*) interaction with the MHC Class II invariant chain (*Cd74*) on B cells, suggestive of a role for the App gene family proteins in modulating immune cell activation and antigen presentation^46,47^ (**Fig. 6E**). Moreover, fibroblast-derived complement factor C3, readily upregulated in cardiac fibroblasts from *T. cruzi*-infected C3H/HeN mice (**Suppl. Fig. 6C**), signalled directly to B-cell complement receptor 2 (*Cr2*/CD21).

**Figure 6.**
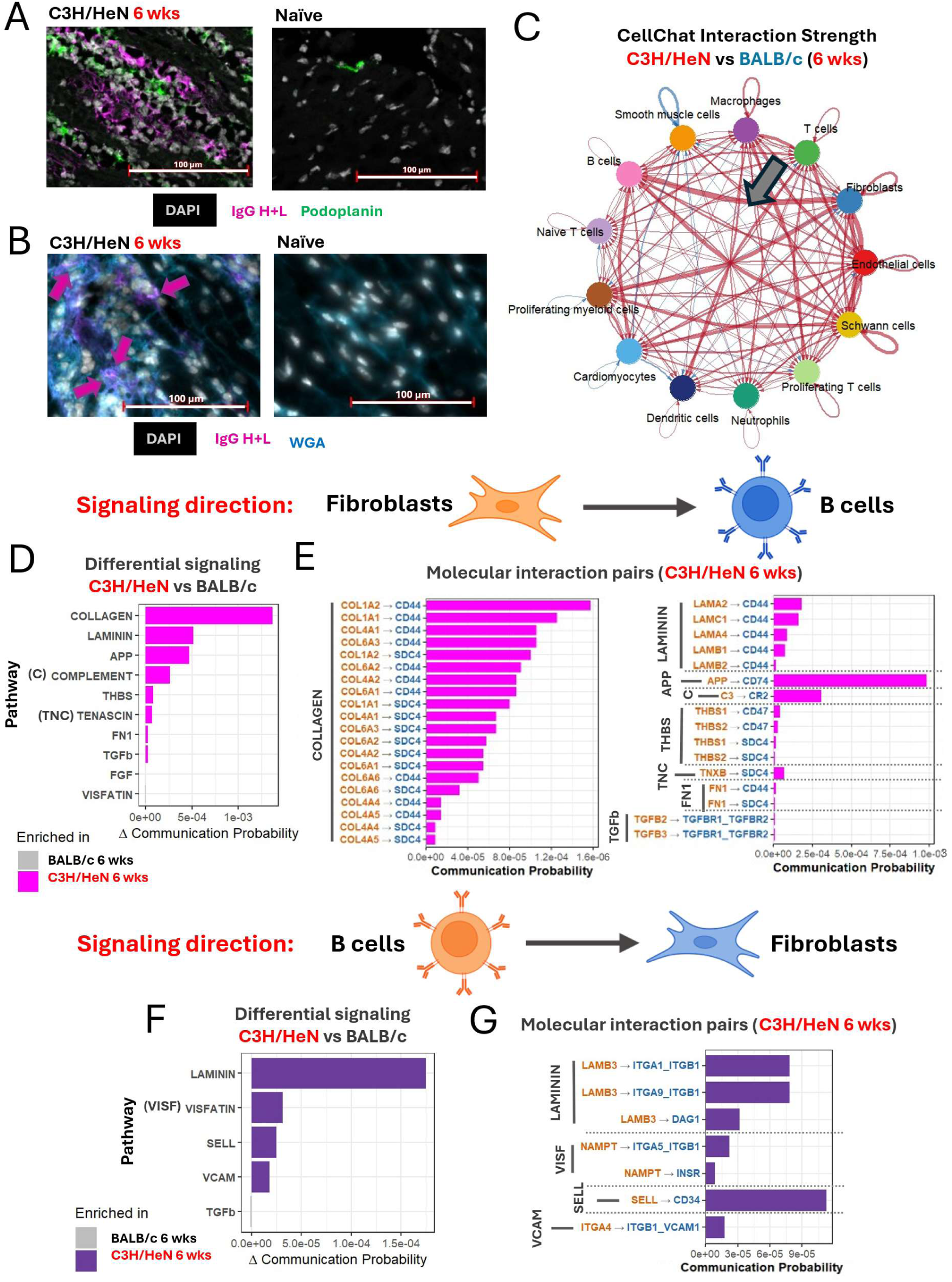
Spatial and computational profiling of fibroblast - B-cell communication during chronic cardiac infection. **A)** Representative immunofluorescence staining demonstrating fibroblasts (Podoplanin) in close spatial proximity to B cells (IgG H+L) in cardiac tissue from C3H/HeN mice at 6 weeks p.i. and naïve age- and strain-matched controls. DAPI marks cell nuclei. **B)** Representative immunofluorescence staining demonstrating increased WGA deposition together with B-cell infiltration (IgG H+L) in cardiac tissue from C3H/HeN mice at 6 weeks p.i. and naïve controls. DAPI marks cell nuclei. **C)** CellChat analysis showing predicted intercellular communication strength between major cardiac cell populations in C3H/HeN and BALB/c mice at 6 weeks p.i. Red indicates interactions enriched in C3H/HeN; blue indicates interactions enriched in BALB/c. The blue arrow highlights the markedly increased predicted interaction between B cells and fibroblasts in C3H/HeN hearts. **D)** Differential signalling pathways identified in the B-cell → fibroblast communication axis between C3H/HeN and BALB/c mice at 6 weeks p.i. **E)** Molecular interaction pairs contributing to pathways upregulated in C3H/HeN mice at 6 weeks p.i. within the B-cell → fibroblast communication axis (orange - signal coming from fibroblasts; blue - signal coming from B cells). **F)** Differential signalling pathways identified in the fibroblast → B-cell communication axis between C3H/HeN and BALB/c mice at 6 weeks p.i. **G)** Molecular interaction pairs contributing to pathways upregulated in C3H/HeN mice at 6 weeks p.i. within the fibroblast → B-cell communication axis (orange - signal coming from B cells; blue - signal coming from fibroblasts).

In the B cell to fibroblast direction, CellChat identified L-selectin (*Sell*), expressed on infiltrating B cells/plasma cells, directly engaging CD34, a sialomucin displayed on cardiac fibroblasts and stromal progenitors^48,49^ (**Fig. 6F-G**). This interaction represents a classical adhesion mechanism driving immune cell tethering, extravasation, and long-term retention within inflamed tissues^49–51^. This was accompanied by an additional classic leukocyte–stromal adhesion pathway, B-cell ITGA4 (*α*4 integrin / VLA-4, paired with *β*1) engaging VCAM1 displayed on fibroblasts (**Fig. 6F-G**). CellChat further identified B-cell-derived visfatin (NAMPT/PBEF (pre-B-cell colony-enhancing factor)) binding *α*5*β*1 integrin (TGA5_ITGB1) or the insulin receptor (INSR) on cardiac fibroblasts (**Fig. 6F-G**). Extracellular NAMPT/visfatin is a well-known pro-inflammatory adipokine/cytokine actively secreted by activated B cells/plasma cells and myeloid cells during chronic inflammation^52^.

CellChat identified an interaction from B cells to fibroblasts via laminin subunit beta-3 (*Lamb3-Itga9/Itgb1*, *Lamb3-Itga1/Itgb1*, and *Lamb3-Dag1*) (**Fig. 6F**-**G**). Single-cell analysis confirmed that *Lamb3* transcript expression was limited to B and T cells from C3H/HeN hearts, low to undetectable in other cell types, and absent in concurrently-sequenced BALB/c samples (**Suppl. Fig. 8 A–B**). Excluding doublets during stringent QC (Materials and Methods) and the absence of expression of other laminin genes in these lymphocytes ruled out the possibility of doublet or ambient RNA contamination (**Suppl. Fig. 8A–B**). The Laminin-332 functional complex requires heterotrimerization of *Lamb3*, *Lama3* and *Lamc2*-gene products^53^. As *Lama3* and *Lamc2* were absent or minimally detected in lymphocytes (**Suppl. Fig. 8A–B**), this expression pattern may reflect an unconventional or non-canonical role for monomeric LAMB3 in immune-stromal communication. Conversely, overall integrin and dystroglycan receptor expression aligned with conventional cell subset expression patterns (**Suppl. Fig. 8C**). *Itga1, Itga9, Igb1* and *Dag1* were expressed in mesenchymal/structural cell populations with *Itga9/Itgab1* additionally expressed on T cells, macrophages and dendritic cells (**Suppl. Fig. 8D**). Immune-cell associated integrins such as *Itga4, Itgal* and *Itgb7* showed expression largely restricted to immune cell populations. The expression of these genes was not strain-specific (**Suppl. Fig. 8D**). Thus, while these computational predictions may reveal an intriguing non-canonical B-cell-fibroblast signalling loop, additional experimentation will be required to define its functional contribution to chronic cardiac pathology.

Collectively, these data show that parasite-reactive and autoreactive antibody-secreting B cells dominate the immune infiltrate and the immune-fibroblast/ECM crosstalk in the fibrosis-prone hearts of C3H/HeN mice during *T. cruzi* infection. Single-cell molecular prediction indicates that B-cell-fibroblast/ECM crosstalk is driven by both extensively characterised canonical (e.g., collagen/laminin-CD44/SDC4) and potentially non-canonical (e.g., LAMB3-Integrins) molecular interactions.

### NKG7^+^CD8^+^ T cells are the dominant source of CCL5 in the heart during chronic Chagas cardiomyopathy

Both natural killer cell granule protein 7 (*Nkg7*), a regulator of lymphocyte granule exocytosis and downstream inflammation in leishmaniasis and malaria^54^, and the chemoattractant CCL5 (*Ccl5*) were among the top upregulated genes in the heart from *T. cruzi*-infected C3H/HeN mice, both at 6 weeks and sustained at 40 weeks p.i. (**Fig. 2A-B**). Cardiac “Th1 cytotoxic” and “NK-like T cell” gene modules were also upregulated and sustained from 6 to 40 weeks p.i. (**Fig. 2C** and **Fig. 4D**).

A deeper 10x Visium spatial transcriptomic map analysis revealed distinct spot-level colocalization of the T-cell receptor subunit *Cd3e*, the CD8 co-receptor *Cd8a*, and *Nkg7* across myocardial sections (**Fig. 7A**). These *Nkg7*^+^ T-cell spots formed widespread niches distributed across both the ventricle and atrium in hearts from infected C3H/HeN mice. In contrast, they were sparse and highly localised in BALB/c hearts (**Fig. 7A**). Spot-level quantification confirmed a striking accumulation of *Cd3e*⁺*Cd8a*⁺*Nkg7*⁺ triple-positive spots in C3H/HeN hearts at 6 weeks p.i. (**Fig. 7B**). To a lesser extent, this spatial signature persisted through late chronic infection in C3H/HeN hearts (40 weeks p.i.) but dropped to baseline levels in BALB/c hearts (**Fig. 7A-B**).

**Figure 7.**
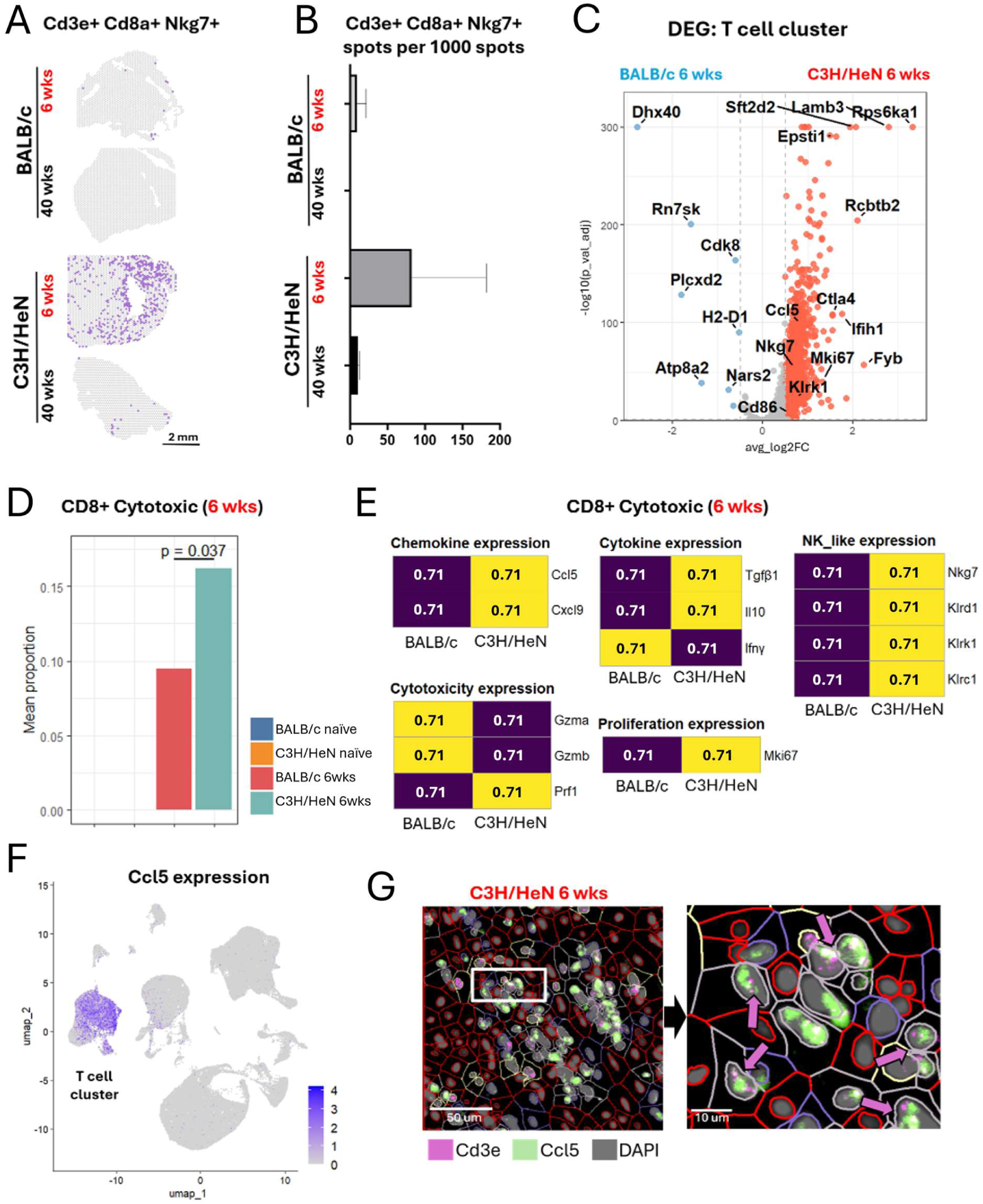
Spatial, transcriptional, and protein-level profiling of cytotoxic T-cell activation during chronic cardiac infection. **A)** Representative spatial transcriptomic maps showing colocalization of Cd3e, Cd8a, and Nkg7 expression across individual tissue spots in C3H/HeN and BALB/c mice at 6 and 40 weeks p.i., highlighting the spatial organisation of cytotoxic T-cell infiltration over time. **B)** Quantification of spot-level colocalization of Cd3e, Cd8a, and Nkg7 expression in C3H/HeN and BALB/c mice at 6 weeks (n = 4 per condition) and 40 weeks (n = 2 per condition). Bars represent mean values with SD. **C)** Volcano plot displaying differential gene expression between C3H/HeN and BALB/c T cells from the scRNA-seq dataset. The top 5 upregulated genes (ranked by log₂ fold-change) from each strain, along with selected additional genes, are labelled. **D)** Mean proportion of cytotoxic Cd8⁺ T cells relative to the total T-cell population across conditions, demonstrating enhanced cytotoxic skewing in C3H/HeN mice during chronic infection. **E)** Modular analysis showing z-scores of selected cytotoxic and activation-associated gene modules derived from scRNA-seq data in C3H/HeN and BALB/c mice at 6 weeks p.i., revealing distinct transcriptional signatures between strains. **F)** Expression of Ccl5 in the scRNA-seq dataset, demonstrating association of Ccl5 expression with the T-cell cluster. **G)** Representative RNAscope images showing co-expression of Cd3e and Ccl5 in cardiac tissue from C3H/HeN mice at 6 weeks p.i., confirming the presence of chemokine-producing T cells within inflamed myocardial regions.

Within the T cell cluster, single-cell differential gene expression revealed a significant upregulation of NK/cytotoxic receptors (*Nkg7*, *Klrk1*), activation markers and regulatory/checkpoint receptors (*Cd86*, *Ctla4*), chemoattractants (*Ccl5*), proliferation factors (*Mki67*), cellular stress/interferon-inducible genes (*Ifih1*, *Epsti1*, *Lamb3*), and signalling adapters (*Fyb*, *Rps6ka1*, *Rcbtb2*) in cardiac C3H/HeN T cells compared to BALB/c T cells during infection (**Fig. 7C**). Both infected BALB/c and C3H/HeN mice exhibited a notable accumulation of cytotoxic CD8⁺ T cells in their hearts; however, the levels were significantly higher in C3H/HeN mice (**Fig. 7D**). Modular analysis highlighted programmes upregulated in C3H/HeN CD8⁺ T cells, including an NK-like signature (*Nkg7*, *Klrd1*, *Klrk1*, *Klrc1*), a chemokine/cytokine signature (*Ccl5*, *Cxcl9*, *Tgfb1*, *Il10*) and a cytotoxicity/proliferation signature (*Prf1* and *Mki67*) (**Fig. 7E**). UMAP feature mapping showed that *Ccl5* expression was strictly restricted to the T-cell cluster, with limit to no expression in other cell types (**Fig. 7F**). Multiplex RNAscope *in situ* hybridization directly confirmed at single-cell spatial resolution co-expression of *Ccl5* within *Cd3e*⁺ T cells in 6-week infected C3H/HeN myocardial tissue (**Fig. 7G**). Thus, orthogonal datasets indicate that NKG7⁺CD8⁺ cytotoxic T cells are the predominant, non-redundant source of CCL5 in the myocardium of *T. cruzi*-infected, fibrosis-prone C3H/HeN mice.

To further dissect heterogeneity of heart-infiltrating T cells during *T. cruzi* infection (**Fig. 4B**), we performed secondary UCell-guided re-clustering (**Suppl. Fig. 9A-D**). Resting naive T-cell pools were excluded from this secondary sub-clustering due to their minimal baseline variation between mouse strains (**Fig. 4B**). While both strains recruited substantial CD4⁺ and CD8⁺ T-cell infiltrates into the myocardium by 6 weeks p.i., single-cell profiling revealed a striking strain-specific dichotomy, particularly so in CD8⁺ T cell differentiation (**Suppl. Fig. 9B**). In hearts from infected BALB/c mice, the CD8⁺ T-cell infiltrate was dominated by conventional αβ cytotoxic CD8⁺ T cells (CD8-Cytox, *Cd8a*⁺/*Cd8b1*⁺/*Gzmb*⁺/*Itga1*⁺/*Cd69*⁺/*Ccl4*⁺/*Nr4a2*⁺), extensively associated with the control of *T. cruzi* infection. γδ T cells were scarce in hearts from this strain (**Suppl. Fig. 9B**). A distinct cardiac *Gzma*^hi^ CD8⁺ T cell population was also detected, this time readily present in both mouse strains (**Suppl. Fig. 9B**), albeit higher in frequency in the heart from infected BALB/c mice (**Suppl. Fig. 9E**). In comparison, the proportion of conventional CD8-Cytox cells was much lower in hearts from infected C3H/HeN mice (**Suppl. Fig. 9E**). In stark contrast, the CD8⁺ T cell infiltrate in hearts from infected C3H/HeN mice was heavily dominated by an unconventional population of NK-like cytotoxic CD8⁺ T cells (NK-like-CD8-Cytox, *Cd8a*⁺/*Klrc1*⁺/*Klrk1*⁺/*Gzmb*⁺/*Gzmk*⁺/*Ccr5*⁺/*Ly6c2*⁺), virtually absent in the hearts from infected BALB/c mice, alongside expanded NKT cells (*Klrd1*⁺/*Klrc3*⁺/*Klrk1*⁺/*Cd244a*⁺) (**Suppl. Fig. 9B** and **E**). Notably, cardiac CD8⁺ T cells, especially NK-like-CD8-Cytox cells, exhibited significant enrichment in γδ T cells (**Suppl. Fig. 9B**).

Thus, chronic infection in fibrosis-prone C3H/HeN hearts seems to shift T-cell differentiation away from conventional protective αβ CD8⁺ T cells toward a dominant, innate-like cytotoxic repertoire composed of unconventional NK-like CD8⁺, NKT, and γδ T cells.

## Discussion

By combining chronic *T. cruzi* infection models with spatial and single-cell orthogonal approaches, here we uncovered an unexpected cellular driver of chronic heart fibrosis: antibody-secreting plasma B cells. These cells represent the most abundant immune population infiltrating the chronically inflamed myocardium and dominate physical and molecular crosstalk with cardiac fibroblasts in hearts from fibrosis-prone *T. cruzi*-infected C3H/HeN mice. We demonstrate a striking accumulation of class-switched plasma B cells (*Fcrl5*^+^, *Jchain*^+^, *Igkc*^+^, *Ighg*^+^) in the fibrotic hearts of chronically *T. cruzi*-infected C3H/HeN mice. Further, our data suggests a previously unnoticed division of labor in the immune system within the chronic cardiac niche, allowing for deeper mechanistic study. Plasma B cells serve as key anchors within the ECM, maintaining localized fibrosis areas. Meanwhile, cytotoxic NKG7^+^ T/NKT cells offer a unique source of the chemoattractant CCL5, aiding in the ongoing recruitment of lymphocytes. Simultaneously, and in agreement with previous data^55–58^, iNOS^+^ M1-skewed macrophages drive CXCL9 and CXCL10 expression to facilitate cytotoxic cell homing. In addition, we detected a persistent cardiac Type I IFN response, previously shown to be detrimental for parasite control^59^, spanning both immune and non-immune structural compartments. This response may further contribute to chronic endothelial activation and pathological tissue remodelling. Moreover, fibroblasts appeared to bridge Type I IFN responses with plasma B cell responses by reacting to IFN signalling and expressing C3, which then engages B cells via complement receptor 2 (CR2). Conversely, chronically-infected BALB/c mice, which show significantly lower cardiac-localised infection and pathology^30^, exhibited a marked cardioprotective phenotype, limiting local lymphocyte retention and cardiac inflammation, and preserving cardiac vascular identity, smooth muscle contractility, and mitochondrial bioenergetic pathways. Importantly, our study provides extensive orthogonal validation at both the transcript and protein levels by using RNAScope and multiplex immunofluorescence in parallel.

In humans, approximately 30% of chronically infected individuals progress to chronic pathology, whereas the other 70% remain asymptomatic despite continued infection^23,60,61^. The clear differences between the *T. cruzi* TcI-JR-infected C3H/HeN and TcVI-CLBR-infected BALB/c mouse models provide a useful experimental route to investigating this clinical spectrum, with the former displaying significantly more disseminated infection, more persistent cardiac parasitism and more severe pathology (heart fibrosis, enteric neurodegeneration)^30,62,63^. Moreover, the bioluminescent reporter *T. cruzi* strains used here allowed us to focus on an early chronic time point (6 weeks p.i.) when parasite burden in the hearts of these different mouse strains is relatively comparable. At this stage, the parasite loads had drastically dropped relative to the acute peak of infection (3 weeks post-infection), showing approximately 100- and 600-fold reductions in overall cardiac parasite burden in BALB/c and C3H/HeN mice, respectively, reaching levels where parasite mRNA was undetectable by 3’ RNA-seq methods in all samples studied. One limitation from this approach is the use of two distinct *T. cruzi* parasite strains across host backgrounds to achieve divergent cardiac pathological outcomes. However, variance partition analysis confirms that host genetic background is the primary driver of cardiac pathology severity in Chagas disease^23,30^. Nonetheless, future work evaluating cross-strain infections will be valuable for fully mapping how nuances in host-parasite interactions impact cardiac immunity and pathology.

Given that available evidence supports Chronic Chagas Cardiomyopathy (CCC) being a T-cell–driven pathology^64–68^, the dominant and widespread accumulation of plasma B cells in the hearts of cardiac fibrosis-prone C3H/HeN mice was unexpected. ELISPOT assays on single-cell cardiac suspensions confirmed that cells infiltrating hearts from chronically-infected mice represented functional antibody-secreting cells with high frequencies of both parasite-reactive and heart-autoreactive IgM^+^ and IgG^+^ B cells. Remarkably, this unusually high local accumulation of antibody-secreting cells and fibrotic scarring occurred despite the extremely low local cardiac parasite burden. Previous reports showed that the spleen of *T. cruzi*-infected mice is the main organ for B cell activation and immunoglobulin production, generating high levels of both parasite-specific and non-parasite-specific (including autoreactive) IgG antibodies, primarily IgG2a^69–72^. In this same model, CD19^neg^CD138^neg^ non-parasite-specific antibody-producing plasma cells, which greatly resemble the phenotype we detected in the heart, can home and persist outside the spleen^73^. Thus, our data support a model of peripheral immune priming and local spatial amplification: in genetically predisposed hosts, minute focal parasite persistence or localised tissue injury in the myocardium acts as a continuous local ‘spark’ that attracts a disproportionately large number of peripherally primed immune effectors, which become physically trapped and maintained within the cardiac matrix. In human chronic Chagas disease, trypanocidal treatment results in a more rapid decline of circulating parasite-specific antibody-secreting B cells compared to circulating anti-*T. cruzi* antibodies, strongly suggesting that long-lived plasma B cells can persist in survival niches years after *T. cruzi* elimination^74^. Given that trypanocidal treatment failed to significantly reduce cardiac clinical deterioration in patients with established Chagas cardiomyopathy, despite significantly reducing parasite load^75^, it becomes paramount to explore whether the myocardial ECM serves as a bona fide survival niche for pathogenic plasma B cells following parasite elimination.

Our study also identified cytotoxic NKG7+ NKT cells as the predominant, non-redundant source of CCL5 in the myocardium of *T. cruzi*-infected, fibrosis-prone C3H/HeN mice. NKG7 expressed by T cells plays critical roles in promoting inflammation during visceral leishmaniasis and malaria^54^, while the CCL5–CCR1/CCR5 axis contributes to both tissue damage and humoral immunity during *T. cruzi* infection^76–79^. Crucially, CCL5 acts directly as a potent B-cell co-activator, promoting antibody-secreting cell generation and enhancing humoral immunity during *T. cruzi* infection^76^; a mechanism that directly underpins local antibody-secreting B cell accumulation and ECM retention observed in the hearts of the chronic C3H/HeN model. The broad significance of CCL5 as a cardiotropic driver is further demonstrated in non-infectious ischemic heart failure, where CCL5 neutralization reduces leukocyte infiltration, matrix metalloproteinase-9 activity, and adverse remodelling^80^. We further discovered a marked expansion of γδ T cells co-expressing *Nkg7*, *Ccl5*, *Cxcl9*, *Il10*, *Ifnγ*, *Mki67* and *Gzmb* in hearts from *T. cruzi*-infected C3H/HeN mice. *T. cruzi* infection in γδ T cell-deficient mice results in significantly lower myocardial inflammation and injury without altering parasite burden^81^. Strikingly, in human chronic Chagas disease, asymptomatic patients maintain γδ T cells with a central-memory phenotype that produce high levels of regulatory IL-10, and this correlates with preserved cardiac function^82,83^. Conversely, in patients with severe Chagas cardiomyopathy, γδ T cells are skewed towards an effector-memory profile enriched for IFN-γ, TNF-α, perforin, granzyme A and CCR5 receptor expression^82,83^. Thus, in cardiac fibrosis-predisposed hosts, *T. cruzi* infection may cause this γδ innate-like T-cell compartment to breach its regulatory IL-10 brake and adopt a heart-homing, pro-inflammatory effector state that contributes to heart tissue remodelling. Moreover, a recent single-cell atlas of human fibrotic diseases shows NKG7^+^CCL5^+^ T cells accumulating in the fibrotic human liver^84^. Thus, beyond the heart, γδ NKG7^+^ NKT/T cells may further represent a pan-organ mechanism of fibrotic tissue remodelling.

The translational significance of our experimental findings is strongly reinforced by a report on single-cell and spatial profiling of human CCC explants, which became available on medRxiv last week^85^. Working completely independently and employing distinct experimental systems [human end-stage cardiac explants and peripheral blood^85^ versus tractable side-by-side mouse models here], the degree of concordance between these investigations and across species is striking. First, in both human end-stage CCC explants and our fibrosis-prone C3H/HeN *T. cruzi* infection model, class-switched antibody-secreting plasma B cells represent the predominant immune population infiltrating the diseased myocardium. Our works complement each other with exceptional synergy regarding local humoral immunity: while targeted BCR sequencing in human CCC hearts reveals clonally expanded B cells sharing unique public clonotypes^85^, functional ELISPOT assays in our model provide direct mechanistic proof that these cardiac plasma B cells actively secrete parasite-reactive and heart-autoreactive IgG and IgM antibodies. Second, in both species, these plasma B cells do not merely circulate but physically embed within the ECM and next to fibroblasts through interactions involving CD44, CD74, syndecan, collagens, and laminins alongside pro-survival stromal crosstalk (BAFF–BCMA / C3-CR2 / Visfatin). Third, this cardiac immune response is accompanied by a distinctive NK/T cell infiltrate. Fourth, both studies highlight the critical role of host background in disease susceptibility, whether reflected in sex-dependent peripheral blood Type I IFN signatures in human progressors or host genetic strain divergence (C3H/HeN versus BALB/c) in mice. That two independent teams, utilizing human clinical explants and mouse models, converged on nearly identical structural and immunological features provides strong validation for the ECM-anchored plasma B cell–fibroblast axis as a conserved, fundamental target in Chagas heart disease.

Beyond its immediate relevance to infectious pathology, our works establish *T. cruzi*-induced cardiomyopathy as a bonafide model for dissecting universal mechanisms of chronic cardiac fibrotic remodelling as well as sterile cardiac autoimmunity. Integrating our findings with recent spatial and single-cell multi-omic human atlases^8,84,86,87^ reveals that intramyocardial plasma B cell retention, direct ECM anchoring, and local autoantibody production are not idiosyncratic features of parasitic infection, but conserved hallmarks of chronic cardiac remodelling across species and disease aetiologies. In the healthy adult human heart, spatial transcriptomics uncovered a constitutive subepicardial immune niche where IgG⁺ and IgA⁺ plasma B cells reside near subepicardial cardiac fibroblasts^8^, proving that cardiac stroma inherently provides a supportive microenvironment for B-lineage cells. Demonstrating the translational relevance of these stromal-immune interactions in non-infectious disease, two very recent transcriptomic studies show significant plasma B cell infiltration alongside fibroblast expansion during chronic cardiac fibrotic remodelling^84,87^. Moreover, integrative profiling of cardiac sarcoidosis, a classic sterile, non-infectious granulomatous inflammatory cardiomyopathy, revealed parallel intramyocardial B-cell expansion, pro-survival stromal cues, and local autoantibody production targeting cardiac structural proteins^86^. Together, through direct ELIPOT mechanistic evidence presented here, BCR repertoire analysis^85^ and antibody cloning from snRNA-seq^86^, our works establish that intramyocardial plasma B cell retention and local autoantibody production represent a hallmark of cardiac fibrotic remodelling across both infectious and sterile aetiologies.

In conclusion, our work addresses a fundamental gap in the broader field of cardioimmunology. Here, we identified a broadly relevant physiological experimental animal model to amplify and study cardiac immune mechanisms implicated in chronic unresolving myocardial fibrosis. Targeting the specific chemoattractant, adhesion, and survival pathways that anchor plasma B cells within the cardiac matrix may offer a broadly applicable therapeutic strategy to halt pathological matrix remodelling in chronic heart disease.

## Materials and Methods

### 1. Mice and *Trypanosoma cruzi* infections

Mouse infections were performed under the UK Home Office project licences PPL PP1572656, 70/8207 and P9AEE04E4 and were approved by the Animal Welfare and Ethical Review Boards of the University of York and London School of Hygiene & Tropical Medicine. All protocols and procedures were conducted according to the UK Animals (Scientific Procedures) Act 1986. Female BALB/c and C3H/HeN mice aged 6-8 weeks were purchased from Charles River (UK) and kept under pathogen-free conditions in individually ventilated cages using a 12h light/dark cycle with constant access to food and water. We used only female mice in this study because previous research with this infection model found no significant differences in infection progression or pathology between males and females^30^. 7-9 weeks old mice were injected intraperitoneally with 10^3^ trypomastigotes from either JR^88^ (DTU I); C3H/HeN) or the CLBR (CL Brener, DTU VI; BALB/c) strains. Both parasite strains were genetically modified to express red-shifted luciferase^30,32^. At experimental endpoints (6 or 40 weeks p.i.), mice were sacrificed by exsanguination under terminal anaesthesia. These mouse/parasite strain combinations were based on previous studies^30^ and aim to reflect the range of cardiac fibrosis outcomes.

### 2. *In vivo* and *ex vivo* imaging (IVIS)

For *in vivo* bioluminescence imaging, infected and control mice were injected with 100 μL of 30mg/mL d-luciferin (in Ca^+2^/Mg^+2^-free DPBS) intraperitoneally, anaesthetised using 2.5% (*v*/*v*) isoflurane in oxygen for 3 min, and then imaged using an IVIS Spectrum system (Perkin Elmer). The detection threshold was determined using uninfected control mice. After imaging, mice were allowed to emerge from anaesthesia on a temperature controlled heat pad and returned to cages. For *ex vivo* imaging, mice were injected with 100 μL of 30mg/mL d-luciferin (in Ca^+2^/Mg^+2^-free DPBS) intraperitoneally, 5-7 min before euthanasia with an overdose of anaesthetic. Trans-cardiac perfusion was performed with 10 mL of DPBS (Dulbecco′s Phosphate Buffered Saline), followed by heart collection. Samples were soaked in DPBS and imaged using an IVIS Spectrum system (Perkin Elmer). To estimate parasite burden in the heart and spleen tissues, regions of interest (ROIs) were drawn using Living Image 4.8.2 to quantify bioluminescence expressed as radiance (photons s^−1^ cm^−2^ sr^−1^). Radiances from equivalent organs/tissues of age-matched, uninfected control mice were measured and the fold change in bioluminescence for each test organ/tissue was calculated.

### 3. Spatial transcriptomics

#### Slide preparation and imaging

Fresh-frozen, OCT-embedded hearts were used to cut 10 µM heart cross-sections directly onto the capture areas of Visium slides (10XGenomics). 24 Visium reactions (6 Visium slides) were carried out.

Methanol fixation, H&E staining and imaging of the Visium slide was performed according to the manufacturer’s (10XGenomics) protocol – “CG000160 Demonstrated Protocol Methanol Fixation and HE Staining”. Imaging was conducted on the Slide Scanner Axioscan Z1 Zeiss instrument at 20x magnification.

#### Library preparation and sequencing

Tissue permeabilization and library construction was performed according to the manufacturer’s (10XGenomics) protocol – “CG000239 Visium Spatial Gene Expression Reagent Kits” user guide. The time for tissue permeabilization was pre-optimised following manufacturer’s instructions. 12 minutes was established as the optimum permeabilisation time for heart tissue and used thereafter on experimental Visium slides. Samples were sequenced by Genewiz/Azenta on Illumina NovaSeq 6000 S4 cell, 10X paired read configuration, dual index with 1% Phi-X spike-in. Samples from C3H/HeN and BALB/c mice were processed and sequenced in parallel.

#### Data processing and analysis

##### Data pre-processing

Fiducial alignment was performed on the HE images of heart cross-sections using Loupe Browser 6.5.0 and JSON files were produced as output. Processing and tissue alignment was conducted via Space Ranger’s count pipeline (Visium, 10x Genomics). Sequencing reads were aligned to a mouse (*Mus musculus*) reference genome (mm10/GRCm38) using the 10x Genomics reference package refdata-gex-mm10-2020-A. Filtered feature matrix files in hdf5 format were produced as one of the outputs and these files were used as input for analysis using Seurat 5.1.0 3, R package. To evaluate local parasite gene expression within cardiac tissue sections, custom reference genomes were generated using 10x Genomics Cell Ranger. FastQ files from C3H/HeN mice infected with *T. cruzi* strain DTU TcI were aligned to the Dm28c reference genome (FASTA and GTF, BioSample ID: SAMN08469708). FastQ files from BALB/c mice infected with *T. cruzi* strain DTU TcVI were aligned to the TCC reference genome (FASTA and GTF, BioSample ID: SAMN08458079). Alignment thresholds and count matrices were evaluated across heart sections from naive and infected mice.

##### QC, filtering and clustering

Cells with at least 300 genes, 300 counts and less than 80% of mitochondrial read counts were used for downstream processing. Normalisation and scaling were performed on each sample using the SCTransform function. The top 2,000 variable features were detected using the FindVariableFeatures function (selection.method = “vst”). The data sets were then integrated by reciprocal PCA integration strategy, fit to integrate large numbers of datasets. Functions such as SelectIntegrationFeatures, PrepSCTIntegration, FindIntegrationAnchors and IntegrateData were used for integration. Following integration, principal components were calculated using RunPCA and the top 20 principal components (supported by ElbowPlot showing decreasing variance beyond the top 20 PCA) were used for computing the ‘k.param’ nearest neighbours for the data sets using the FindNeighbours function (k.param = 20). Following, FindClusters function was used for semi-unsupervised clustering (resolution = 0.5) and UMAP reduction was performed. Clusters were then manually annotated by marker gene identification (identifying marker genes that are uniquely or highly expressed in each cluster) with Enrichr web utility (https://maayanlab.cloud/Enrichr/enrich).

##### Mapping gene expression on spatial plots

The mapping of gene expression of selected genes on the corresponding tissue sections was conducted using Loupe Browser 6.5.0 and cloupe files (generated during data processing via Space Ranger) were used as input.

##### Differential gene expression analysis (DEG)

Detection of differentially expressed genes between experimental conditions (time points p.i. and mouse strain comparisons) was performed using the FindMarkers function, specified to return both upregulated as well as downregulated genes. The DEG table was further filtered by Log_2_FC > 0.5 and P adjusted values < 0.05 (P values adjusted by the default Bonferroni correction method). For all DEG calculations the “SCT” assay and “data” slot were utilised and performed by the default Wilcoxon rank-sum method. Data was visualised using volcano plots (ggplot2; ggrepel packages).

##### Gene ontology (GO)

GO analysis was conducted on the integrated object using the ClusterProfiler R package. Filtered (by Log_2_FC > 0.5 and P adjusted values < 0.05) DEG table of the “C3H/HeN 6 weeks p.i. vs BALB/c 6 weeks p.i.” comparison was used as input to identify significant biological processes upregulated in these conditions. Top 30 biological processes (ranked by adjusted p-value) identified through GO analysis were visualised.

##### SPOTlight cell type deconvolution

SPOTlight was used to predict the proportional abundance of distinct cardiac cell types across spatial transcriptomic tissue sections. A subset of our annotated scRNA-seq dataset (Methods Section 5) served as the single-cell reference. Marker genes for each cell type were identified from the scRNA-seq differential expression analysis, and the top weighted markers per cluster were selected to construct the SPOTlight marker gene table. The scRNA-seq reference object was converted to a SingleCellExperiment format, and spatial transcriptomic data were extracted as raw count matrices. SPOTlight was then run using the single-cell counts, spatial counts, cluster assignments, and curated marker gene table. The method applies non-negative matrix factorisation (NMF) to learn cell-type-specific topic profiles and subsequently performs seeded non-negative least squares regression to estimate cell-type proportions for each spatial spot. The resulting deconvolution matrix contained predicted proportions for all annotated cell types across the tissue section. Topic profiles and interaction metrics were inspected to confirm model stability and biological coherence. Spatial visualisation of cell-type proportions was performed using scatter-pie overlays on the tissue coordinates. Cell-type proportion numbers were extracted and plotted as bar charts.

##### Gene Module Scoring

Gene-expression modules were constructed to summarise coordinated transcriptional programs across immune, stromal, endothelial, and metabolic cell populations. Gene modules (Table 1.) representing key biological processes were defined based on established marker genes. For each sample, raw counts were normalised and averaged per condition. Module scores were then computed by calculating the mean expression of all genes within each module across the averaged expression matrix. Only genes present in the dataset were included in each calculation. Resulting module-by-condition matrices were z-scaled to enable comparison of relative module activity across experimental groups. Heatmaps were generated to visualise module activity patterns across naïve and infected conditions at both time points, allowing identification of strain-specific and infection-associated transcriptional programs.

**Table 1.** Gene modules used for module score calculation and biological annotation of cell states.

| Module | Description | Genes |
|---|---|---|
| **Plasma_Cell_Response** | Antibody-secreting plasma cell program marked by immunoglobulin production, plasma cell differentiation and B-cell survival signals. | *Cxcl13, Jchain, Ighg1, Ighg2c, Ighg2b, Ighg3, Igkc, Tnfsf13b, Iglc1, Mzb1* |
| **Naive_B_Cell_Identity** | Resting B-cell program characterized by B-cell receptor expression and antigen-responsive signaling machinery. | *Ighm, Ighd, Cd79a, Cd79b, Cd19, Ms4a1, Cd72, Cd40, Cd84* |
| **Th1_Cytotoxic_Effector** | Cytotoxic type-1 T-cell program associated with IFN-γ production, T-cell activation and target cell killing. | *Tbx21, Ifng, Cd3d, Cd3e, Cd2, Cd96, Cd160, Prf1, Gzmb, Gzmk, Gzma, Ctsw, Cxcr6, Ccl5, Il21r, Zap70, Lck, Lat* |
| **NK_Cytotoxic_Program** | Innate-like cytotoxic lymphocyte program enriched for NK-cell receptors and effector functions. | *Klrd1, Klrk1, Klrb1c, Ncr1, Nkg7, Prf1, Ctsw, Cst7, Il18rap, Tyrobp, Trem2* |
| **Inflammatory_Macrophages** | Inflammatory macrophage program characterized by pathogen sensing, cytokine signaling and innate immune activation. | *Nos2, Acod1, Nod2, Tlr2, Tlr12, Tlr13, Ccl2, Ccl4l2, Ccl7, Il1rn, Tnfaip3, Msr1, Fcgr1, Clec4e* |
| **Type_I_IFN_Response** | Type I interferon response program enriched for antiviral and interferon-stimulated genes. | *Ifit1, Ifit2, Ifit3, Oasl1, Oasl2, Irf7, Irf8, Zbp1, Gbp5, Rsad2* |
| **Inflammatory_Cytokines** | Pro-inflammatory cytokine and chemokine program involved in immune cell recruitment and activation. | *Tnf, Ccl2, Ccl4, Ccl5, Ccl7, Ccl8, Cxcl9, Cxcl10, Cxcl16, Il18bp* |
| **Fibrotic_Remodeling** | Extracellular matrix remodeling and fibrotic tissue repair program associated with collagen deposition and matrix turnover. | *Col1a1, Col1a2, Fn1, Thbs1, Lox, Loxl1, Mmp12, Mmp14, Timp1, Tgfbi* |
| **Activated_Endothelium** | Activated endothelial cell program associated with leukocyte adhesion, vascular inflammation and angiogenic remodeling. | *Vcam1, Icam1, Edn1, Vegfd, Plvap, Pdgfc, Itgav, Itga5, Itgb3, Vamp8* |
| **Arterial_Vascular_Identity** | Gene program defining mature vascular and arterial endothelial cell identity. | *Dll4, Efnb2, Ednra, Sparcl1, Col4a5, Cavin2, Pcdh12, Thsd1, Tmem204, Scube2* |
| **Contractile_Smooth_Muscle** | Contractile smooth muscle program involved in vascular tone regulation and vessel function. | *Acta2, Cnn1, Lmod1, Myzap, Mylk4, Kcnj2, Kcnd2, Rgs4, Aqp1, Aqp7* |
| **Lipid_Metabolism** | Lipid metabolic and energy homeostasis program associated with fatty acid utilization and redox balance. | *Pdk4, Elovl6, Scd1, Scd4, Thrsp, Adipoq, G0s2, Txnrd3, Hmgcl, Acox1, Acot1, Acot2, Nnt, Etfrf1* |
| **Secretory_Epithelial_Identity** | Secretory epithelial cell program associated with barrier maintenance, fluid transport and protective factor secretion. | *Scgb1a1, Scgb1c1, Bpifa1, Car4, Aqp6, Pla2g2d, Upk3b, Lgals4, Hnmt, Inmt* |
| **Stromal_Identity** | Stromal cell identity program enriched for developmental and differentiation-associated transcription factors. | *Klf4, Klf9, Klf15, Meox2, Casz1, Runx1t1, Ebf2, Ebf3, Nr2f1, Ctf1, Ccn3, Ccn5* |
| **Stromal_Quiescence** | Differentiated cell maintenance program associated with transcriptional stability and cellular homeostasis. | *Klf4, Klf9, Klf15, Meox2, Casz1, Runx1t1, Ebf2, Ebf3, Nr2f1, Ctf1, Tshz2, Foxk1* |
| **Structural_ECM** | Extracellular matrix structural program associated with matrix organization and tissue support. | *Fn1, Col4a5, Sparcl1, Ccn3, Ccn5, Timp4, Colgalt2, Pdgfc* |
| **Vascular_Communication** | Vascular signaling program involved in endothelial communication, vessel patterning and homeostasis. | *Dll4, Efnb2, Edn1, Ednra, Vegfd, Pdgfc, Cavin2, Thsd1* |

### 4. RNAScope^TM^ Assay

Fresh-frozen, OCT embedded heart tissues were used to cut 10 µM heart cross-sections on super-frost slides. The RNAScope^TM^ Assay manufacturer’s protocol “RNAscope^TM^ HiPlex12 Reagent Kit (488, 550, 650, 750) v2 Standard Assay” was followed for sample pretreatment and preparation of fresh frozen samples. Three rounds of detection were carried out and included markers and controls present in Table 1.

Imaging was carried out on the Slide Scanner Axioscan Z1 Zeiss instrument at 20x magnification. Identical exposure times and threshold settings were used for each channel on all sections of similar experiments. Images from rounds 1, 2 and 3 from each sample were registered together using the Hiplex Image Tissue Registration Software (Biotechne) and all subsequent analysis on the registered image was conducted in QuPath. Cell segmentation was carried out using StarDist (QuPath extension) and the automated segmentation was based on the DAPI nuclei staining. Cell counts of cells expressing a single marker or co-expressing selected double markers was based on the presence or absence of this marker/s in the segmented cells.

### 5. 18-plex ChipCytometryTM assay

Fresh-frozen, OCT embedded heart tissues were used to cut 10 µM heart cross-sections on PRSM coverslips (Canopy Biosciences). The coverslips containing the tissue sections were then assembled in the chips following the manufacturers protocol and the PRSM storage buffer was added. Assembled chips were shipped to Canopy Biosciences (Germany) for staining and imaging.

The assay panel included 18 target markers: **CD45, CD19, FoxP3, CD4, CD8a, Ki67, CD3, CD335, CD11b, CD11c, MHCII, CD69, IgG/R, F4/80, Gr1, PanCK, Cp38/Podoplanin, DNA** (Table 2) and these markers were used to immunophenotype chosen cell types on the heart sections, as presented in Table 3.

**Table 2.** Marker genes. Each round allowed for staining with 4 cell type/chemokine marker genes. Additionally, each round included DAPI staining to mark the nuclei.

| Staining round | Cell type/chemokine marker gene | Cell type/chemokine represented by the marker gene | Fluorophore |
|---|---|---|---|
| Round 1 | Ttn | Cardiomyocytes | AF 488 |
|  | Col1a1 | Fibroblasts | ATTO 550 |
|  | Acta2 | Smooth muscle cells | ATTO 647N |
|  | Tgfb1 | Tgf-B | AF 750 |
| Round 2 | Nrp3 | Endocardial cells | AF 488 |
|  | Ighm | B cells | ATTO 550 |
|  | Ighg1 | B cells | ATTO 647N |
|  | Sdc1 | Plasma B cells | AF 750 |
| Round 3 | Nr4a1 | B cell activation | AF 488 |
|  | Fcgr1 | Macrophages | ATTO 550 |
|  | Cd3e | T cells | ATTO 647N |
|  | Ccl5 | Ccl5 | AF 750 |
| All rounds | DAPI | Nuclei marker | - |
| Positive control | Ppib | Housekeeping gene | - |
| Negative control | dapB | *Escherichia coli* gene | - |

**Table 3.** 18-plex ChipCytometryTM assay staining panel.

| Cycle | Epitope | Clone | Fluorophore | Compartment | Dilution | Vendor |
|---|---|---|---|---|---|---|
| 1 | CD45 | 30-F11 | FITC | Membrane | 1500 | Biolegend |
|  | CD19 | 1D3 | PerCP-Cy5.5 | Membrane | 50 | Biolegend |
|  | FoxP3 | FJK-16S | PE | Nuclear | 50 | Thermo Fisher Scientific |
| 2 | CD4 | RM4-5 | PerCP-Cy5.5 | Membrane | 300 | Biolegend |
|  | CD8a | 53-6.7 | BUV395 | Membrane | 100 | BD |
|  | Ki67 | SolA15 | PE | Nuclear | 50 | Thermo Fisher Scientific |
| 3 | CD3 | 17A2 | PerCP-Cy5.5 | Membrane | 200 | Biolegend |
|  | CD335 | 29A1.4 | PE | Membrane | 50 | Biolegend |
| 4 | CD11b | M1/70 | PerCP-Cy5.5 | Membrane | 300 | Biolegend |
|  | MHCII | M5/114.15.2 | FITC | Membrane | 600 | Biolegend |
|  | CD69 | H1.2F3 | PE | Membrane | 100 | Biolegend |
| 5 | IgG-PE | poly | PE | Membrane | 1500 | Thermo Fisher Scientific |
| 5b | mouse IgG isotype blocking | -- | none | NA | 300 | Thermo Fisher Scientific |
| 6 | CD11c | N418 | PE | Membrane | 50 | Biolegend |
|  | F4/80 | BM8 | PerCP-Cy5.5 | Membrane | 100 | Biolegend |
|  | Gr1 | RB6-8C5 | BUV395 | Membrane | 500 | BD |
|  | panCK | C11 | AF488 | Membrane | 1200 | Biolegend |
| 7 | anti-AF488 | -- | Quench | NA | 100 | Thermo Fisher Scientific |
| 8 | Gp38/Podoplanin | 8.1.1 | AF488 | Membrane | 100 | Thermo Fisher Scientific |
|  | DNA | -- | Hoechst 33342 (BV/BUV channel) | Nuclear | 100k | Invitrogen |
| 9 | anti-AF488 | -- | Quench | NA | 100 | Thermo Fisher Scientific |
| 10 | Helix Green (DNA) | -- | AF488 | Nuclear | 5 Mio | Biolegend |
|  | PI (DNA) | -- | PE | Nuclear | 50k | Hölzel Diagnostika |

**Table 4.** Characterization of immune cells. Cells were characterized depending on the established marker combinations.

| Phenotype | Marker combination |
|---|---|
| All cells | DNA+ |
| Leukocytes | CD45+ |
| T cells | CD45+, CD3+, CD335- |
| CD4+ T cells | CD45+, CD3+, CD335-, CD4+, CD8- |
| CD8 + T cells | CD45+, CD3+, CD335-, CD4-, CD8+ |
| Activated T cells | CD45+, CD3+, CD335-, MHC II+ |
| T regulatory cells | CD45+ CD3+ CD4+ FoxP3+ |
| B cells | CD45+, CD3-, CD335-, IgG/R+ |
| CD19 B cells | CD45+, CD3-, CD335-, CD19+ |
| Dendritic cells | CD45+, CD3-, CD335-, CD11c+, F4/80- |
| Monocytes/Macrophages | CD45+, CD3-, CD335-, CD11c-, F4/80+ |
| Neutrophils | CD45+, CD3-, CD335-, CD11c-, F4/80-, Gr1+,CD11b+ |
| Proliferating immune cells | CD45+, Ki-67+ |
| Fibroblasts | CD45-, Gp38+ |

Thresholding, cell segmentation, and quantification of target cell populations were performed using QuPath. Quantitative data (Fig. 3.) were visualized and plotted in GraphPad Prism 9 (GraphPad Software), and statistical comparisons between groups (BALB/C 6wks vs C3H/HeN 6wks) were performed using two-tailed unpaired Student’s *t*-tests.

### 6. scRNA-seq

#### Preparation of heart single cell suspensions

The heart tissue dissociation included both enzymatic digestion and mechanical dissociation. The Multi tissue dissociation kit 2 (Miltenyi Biotec) was used in combination with the gentleMASC dissociator (Miltenyi Biotec) following manufacturer’s protocol - “Dissociation of adult mouse heart using the Multi Tissue Dissociation Kit 2”. After tissue dissociation, debris was removed with Debris Removal Solution (Miltenyi Biotec) following manufacturer’s instructions, with reagents adjusted to small tissue quantities (<100 mg). Red blood cells were then removed with ACK Lysing Buffer (ThermoFisher) – for this, cell pellets were resuspended in 1 ml of ACK Lysis Buffer, incubated 2-5 min. at room temperature and resuspended in 10 ml of 20% FBS DMEM was. Cells were then centrifuged at 300 g, resuspended in 1 ml of PBS and counted using trypan blue and a haemocytometer. Cell counts in each sample ranged from 150,000 cells to 600, 000 cells.

#### Library preparation and sequencing

Cells obtained above were fixed and stored at −80°C for less than six months before library preparation following Evercode WT Mini V3 or Evercode WT V3 protocols (Parse Biosciences). Samples were sequenced by Genewiz/Azenta on an Illumina NovaSeq X Plus 25B flow cell.

#### Data processing and analysis

##### Data pre-processing

Data processing of raw counts in the form of fastq files was conducted through the Trailmaker (Parse Biosciences) software and a mouse (*Mus musculus*) reference genome set was used.

##### Doublets exclusion

Doublets were identified and removed using scDblFinder, a model-based approach for doublet detection in scRNA-seq data. Raw, per-sample Seurat objects were first converted into SingleCellExperiment objects, and scDblFinder was applied independently to each sample prior to any normalisation or downstream processing. For every cell, scDblFinder generated a doublet probability score and a binary doublet classification. These annotations were transferred back to the corresponding Seurat objects, and cells classified as doublets were excluded. Following doublet removal, singlet-only Seurat objects were generated for each sample and saved for downstream analysis. The number and proportion of removed cells were recorded for all samples to ensure consistent doublet rates across conditions. All subsequent analyses, including normalisation, clustering, differential expression, and integration, were performed exclusively on the singlet-filtered datasets.

##### Quality control and filtering

Cells with at least 300 genes, counts between 300 and 25,000 as well as less than 60% of mitochondrial read percentage were used for downstream processing.

##### Normalisation, data integration and UMAP clustering

Normalisation and scaling were performed on each sample using the SCTransform function. The top 5,000 variable features were detected using the FindVariableFeatures function (selection.method = “vst”). The data sets were then integrated by reciprocal PCA integration strategy, fit to integrate large numbers of datasets. Functions such as SelectIntegrationFeatures, PrepSCTIntegration, FindIntegrationAnchors and IntegrateData were used for integration. Following integration, principal components were calculated using RunPCA and the top 30 principal components (supported by ElbowPlot showing decreasing variance beyond the top 30 PCA) were used for computing the ‘k.param’ nearest neighbours for the data sets using the FindNeighbours function. Following, FindClusters function was used for semi-unsupervised clustering (resolution = 0.3) and UMAP reduction (dims = 30) was performed. Clusters were then manually annotated by marker gene identification (identifying marker genes that are uniquely or highly expressed in each cluster) with the use of Enrichr web utility (https://maayanlab.cloud/Enrichr/enrich).

##### Differential gene expression analysis (DEG)

Detection of differentially expressed genes between experimental conditions (time points p.i. and mouse strain comparisons) was performed using the FindMarkers function, specified to return both upregulated as well as downregulated genes. The DEG table was further filtered by Log_2_FC > 0.5 and P adjusted values < 0.05 (P values adjusted by the default Bonferroni correction method). For all DEG calculations the “SCT” assay and “data” slot were utilised and performed by the default Wilcoxon rank-sum method. Data was visualised using volcano plots (ggplot2; ggrepel packages). Additionally, heatmaps were generated based on the conducted DEG analysis for the following subclusters: B cells, T cells, Macrophages, Dendritic cells (DCs), Neutrophils, Fibroblasts, Endothelial cells and Smooth muscle cells (SMCs), each showing the top 25 highest z-scores in these cell types for C3H/HeN at 6 weeks p.i. (Fig. 4.) and for BALB/c at 6 weeks p.i. (Suppl Fig. 4.) compared with other conditions. The 25 genes were selected from the top 50 upregulated genes in C3H/HeN 6 weeks compared to C3H/HeN naïve or in BALB/c 6 weeks compared to BALB/c naïve based on z-score ranking after calculating the average expression of each gene across all conditions.

##### Gene ontology (GO) for Macrophage cluster

Gene ontology analysis was conducted on the subsetted macrophage cluster using the ClusterProfiler R package. Filtered (by Log_2_FC > 0.5 and P adjusted values < 0.05) DEG tables (“C3H/HeN 6 weeks p.i. vs C3H/HeN naïve” or “BALB/c 6 weeks p.i. vs BALB/c naïve”) were used as input to identify significant biological processes upregulated in C3H/HeN mice 6 weeks p.i. or BALB/c mice 6 weeks p.i. Top 15 biological processes (ranked by adjusted p-value) identified through GO analysis were visualised for each analysed condition.

##### Cell–Cell Communication Analysis (CellChat)

Cell–cell communication was analysed using CellChat. For every condition, the corresponding singlet-filtered Seurat object was subsetted and converted into a CellChat object using the annotated cell-type identities as grouping labels. The CellChatDB.mouse ligand–receptor database was used as the reference interaction framework. Each dataset underwent the standard CellChat preprocessing workflow, including identification of over-expressed genes and interactions, and projection of expression data onto the mouse protein–protein interaction network. Pathway-level communication scores were then calculated. To compare communication patterns between strains, CellChat objects from C3H/HeN 6 weeks p.i. and BALB/c 6 weeks p.i. were merged and the differences in interaction count and/or strength were visualised with the netVisual_diffInteraction plot. Following, to compare intercellular communication between B cells and fibroblasts across mouse strains, we performed directional signalling analyses. For each condition, we extracted all ligand-receptor interactions involving B cells as senders and fibroblasts as receivers, and separately all interactions involving fibroblasts as senders and B cells as receivers. For each signalling direction, interaction probabilities were summed at the pathway level to obtain a total communication score per pathway. Pathway-level scores from BALB/c and C3H/HeN mice were then merged, and a differential communication metric (Δ probability) was calculated as the difference between strains. Pathways were ranked according to this Δ value to identify signalling routes enriched in either BALB/c or C3H/HeN hearts. To visualise these differences, pathway-level Δ communication probabilities were plotted using bar charts with pathways ordered by effect size. For the pathways that emerged as showing strong strain-specific enrichment during this analysis, we further examined individual ligand-receptor pairs contributing to the pathway. For each pathway, all constituent ligand-receptor interactions were extracted for the relevant signalling direction (B cell → Fibroblast or Fibroblast → B cell), and interaction probabilities were plotted to highlight the dominant ligand-receptor pairs driving the communication signal.

##### Macrophage, Fibroblast and T cell population sub-clustering and annotation

Macrophages/Fibroblasts or T cells were extracted from the integrated dataset and subjected to secondary unsupervised clustering analysis. Cells were dimensionally reduced using PCA and the number of principal components retained for downstream analyses was determined using PCA variance distribution (Elbow Plot) and biological interpretability of the resulting clustering structure. A shared nearest neighbor graph was constructed from the selected principal components, followed by graph-based clustering and UMAP visualization to identify transcriptionally distinct Macrophages, Fibroblasts or T cells subpopulations. To characterize each Macrophages, Fibroblasts or T cells subcluster, cluster-specific marker genes were identified using Seurat’s FindMarkers function and the Top25(for Macrophages) and Top50 (for Fibroblasts and T cells) DEG genes were visualized by heatmap. From the Top25 or Top50 DEG genes, candidate marker genes were selected based on cluster specificity assessed by FeaturePlots and biological relevance determined through Enrichr pathway enrichment analysis. These genes were used to calculate UCell enrichment scores for each subcluster (Table 5.) and were visualized on UMAP embeddings using FeaturePlots, enabling assessment of the spatial distribution of each transcriptional program across the Macrophages, Fibroblasts or T cell compartment. These UCell scores were subsequently used to support biological annotation of subclusters.

**Table 5.** Macrophage, Fibroblast and T cell subclusters with the genes selected from the Top25 (Macrophage) or Top50 (Fibroblasts and T cells) DEG genes for UCell enrichment score calculations.

| **Cell type subset abbreviation** | **Cell type subset full name** | **Cell type** | **Genes selected for UCell enrichment scores** |
|---|---|---|---|
| Comp. Resident | Complement resident | Macrophage | *Cfp, Pltp, Lacc1, Atp8b4, Mgl2* |
| Resident 1 | Resident 1 | Macrophage | *Nr3c2, Cpne8, Magi1, Samd4, Adam22, Ppp1r9a* |
| Resident 2 | Resident 2 | Macrophage | *Cd163, Hpgd, Slco2b1, Prkg1, Clmp* |
| Comp. Inflamm. | Complement Inflammatory | Macrophage | *Cxcl9, Acod1, Ly6c2, C3, Cfb, Clec4e* |
| Inflamm. | Inflammatory | Macrophage | *Trem1, Trem3, Plac8, Sell, Hp, Ace* |
| Regenerative | Regenerative | Macrophage | *Tgfb3, Ptgs1, Acp5, Id2, Id3* |
| Proliferative | Proliferative | Macrophage | *Kif18b, Kif14, Kif4, Ccna2, Pclaf, Neil3, Nuf2, Hmmr* |
| F-Base-Int | Baseline Interstitial Fibroblasts | Fibroblast | *Polg2, Ppp1r10, Hsph1, Ccn1, Mat2a, Jup, Junb, Ptger4, Col5a3, Spry1, Atf3, Hes1, Btg2, Klf2* |
| F-C3-IFN | Complement (C3) Interferon Fibroblasts | Fibroblast | *Gabbr1, Mt2, Spon1, Ror1, Hmcn2, Cacna1c, Fkbp5, Ly6c1, Mt1, AU020206, C3, Gbp2b, Gbp7, Gbp9, Nlrc5, Trim30a* |
| F-Activ | Activated Fibroblasts | Fibroblast | *Gfra1, Rn7sk, Rgs7, Dpep1, Grik2, Pde4b, Hlf, Thsd7a, Dbp, Slit3, Il1r1, Cxcl1, Osmr, Irf1, Iigp1, Mme* |
| F-Base-Str | Baseline Structural Fibroblasts | Fibroblast | *Nkain3, Piezo2, Smad6, Igfbp3, Eya4, Pdlim3, Olfml2b, Hmcn1, Apoe, Hecw2, Eln, Fbln1, Fbln5, Col14a1, Sulf2, Slit2* |
| F-WntX | WntX Fibroblasts | Fibroblast | *Erbb4, Pdzrn4, Rerg, Cadm1, Nrxn1, Kcnma1, Kcnq5, Rspo3, Wif1, Wnt5a, Frzb, Axin2, Prg4, Clu, Rgs5, Col12a1* |
| CD8-Cytox | CD8 Cytotoxic | T cells | *Cd8a, Cd8b1, Itga1, Cd69, Ccl4, Il21r, Tnfrsf1b, Nr4a2, Dusp2, Cd226* |
| Activ-CD4 | Activated CD4 | T cells | *Cd4, Cd40lg, Icos, Ctla4, Cd28, Maf, Tox, Nfatc1, Cd44, Slamf1* |
| NK-like-CD8-Cytox | NKT CD8 Cytotoxic | T cells | *Cd8a, Cd8b1, Klrc1, Klrk1, Gzmb, Gzmk, Ccr5, Ly6c2, Itga1, Ptpre* |
| Gzma-<sup>hi</sup>- CD8 Cytox | Gzma high CD8 Cytotoxic | T cells | *Gzma, Cd8a, Cd8b1* |
| Prolif T cells | Proliferating T cells | T cells | *Mki67, Top2a, Birc5, Ccna2, Foxm1, Tpx2, Cenpf, Nusap1, Uhrf1, Cdc6* |
| NKT cells | NK-like cells | T cells | *Klrd1, Klrk1, Klrc3, Klrb1f, Cd244a, Itgax, Tox, Cd200r1, Cd200r4, Farp1* |
| NK cells | Natural killer cells | T cells | *Eomes, Tyrobp, Syk, Itgam, Irf8* |
| Il1rl1-Pos T reg | Il1rl1 Positive regulatory T cells | T cells | *Il1rl1, Pparg, Irs2* |

##### Macrophage, Fibroblast and T cell subcluster abundance analysis

To assess changes in subpopulation abundance between experimental groups, the proportion of cells assigned to each sub-cluster was calculated relative to the total number of macrophages, fibroblasts or T cells within each biological replicate. For each replicate, cluster abundance was defined as: Cluster proportion = Number of cells in sub-cluster/Total number of cells (macrophages, fibroblasts or T cells). Mean cluster proportions were subsequently calculated for each experimental group (BALB/c naïve, C3H/HeN naïve, BALB/c 6 wks p.i., and C3H/HeN 6 wks p.i.) and visualized as bar plots. Statistical comparisons were restricted to the infected groups, as these represented the biologically relevant comparison and were available as independent biological replicates. Differences in cluster proportions between BALB/c and C3H/HeN mice at 6 wks p.i. were assessed using two-sided Wilcoxon rank-sum tests performed on replicate-level cluster proportions. Raw *P* values were reported and displayed on the corresponding plots.

##### Identification and Quantification of γδ T Cells

γδ T cells were identified from the T-cell subset based on expression of canonical γδ T-cell receptor (TCR) genes. Cells expressing either ***Trdc*** or ***Trgc2*** were classified as γδ T cells. The resulting γδ T-cell annotation was stored as metadata for downstream analyses. To compare γδ T-cell abundance between infected mouse strains, analyses were restricted to cells originating from infected samples (BALBc 6wks and C3HHeN 6wks). For each biological replicate, the proportion of γδ T cells was calculated as: Proportion of γδ T cells = Number of γδ T cells/ Total classified T cells. Proportions were first calculated at the individual sample level and subsequently averaged across biological replicates within each experimental group. Differences in γδ T-cell proportions between BALBc 6wks and C3HHeN 6wks mice were assessed using a two-sided Wilcoxon rank-sum test. Group means were visualized as bar plots, and the corresponding p-value was displayed on the Fig. γδ T cells were highlighted on the UMAP by overlaying their coordinates onto cluster-based dimensional reduction plots. Separate UMAPs were generated for each experimental condition using split-panel visualization to assess the spatial distribution of γδ T cells across samples.

### 7. Immunofluorescence staining and imaging

Two immunofluorescence staining protocols were performed on fresh frozen cardiac tissue sections (10 µm) to assess B-cell infiltration and fibroblast-rich or fibrotic regions.

#### IgG H+L, Podoplanin, DAPI

Sections were fixed for 5 min in cold methanol at RT, washed in PBS, and blocked with 1% BSA (Gibco, Thermo Fisher Scientific). Slides were incubated with AF555-conjugated F(ab’)2-Goat anti-Mouse IgG (H+L) (1:100, Invitrogen) and AF488-conjugated anti-mouse Podoplanin (1:100, Biolegend) for 1 h at RT, washed, and subsequently counterstained with DAPI. Sections were washed, mounted, and imaged on a ZEISS LSM 780 Confocal Microscope using 20× magnification.

#### IgG H+L, WGA, DAPI

Sections were fixed for 5 min in cold methanol at RT, washed in PBS, and incubated with wheat germ agglutinin (WGA) (1:1000 Invitrogen) for 2 h at RT. After PBS washing, sections were blocked with 1% BSA (Gibco, Thermo Fisher Scientific). Slides were then stained with AF555-conjugated F(ab’)2-Goat anti-Mouse IgG (H+L) (1:100, Invitrogen) for 1 h, washed, and counterstained with DAPI. Sections were washed, mounted, and imaged on a ZEISS Axio Observer 7 using 40× magnification.

### 8. B-cell ELISpot Assay

#### Plate preparation and coating

Plates were activated with 100 µL of 100% ethanol for 1 min, washed three times with PBS, and coated either with an in-house produced crude freeze-thaw protein extraction of tissue cultured *T. cruzi* trypomastigotes (200 µg/mL) or a high-quality microbiologically graded bovine heart extract (200 µg/mL, Sigma-Aldrich) to detect antigen-specific ASCs. Anti-mouse IgM and IgG capture antibodies (10 µg/mL, SouthernBiotech) to measure total IgM and IgG-producing ASCs. Coated plates were incubated overnight at 4 °C.

#### Blocking and cell incubation

Plates were washed with PBS and blocked for 2 h at 37 °C using RPMI with 10% FBS. Single-cell suspensions from mouse hearts were added in complete RPMI (RPMI + 10% FBS, 1% glutamine, 1% Penicillin-Streptomycin, 0.1% 2-mercaptoethanol) and incubated overnight at 37 °C (≤20 h).

#### Detection and development

Cells were removed and plates washed with 0.05% PBS-Tween. Anti-Ig-biotin (1:1000) (SouthernBiotech) was added for 1 h at room temperature, followed by alkaline phosphatase–extravidin (1:5000) for 30 min. Spots were developed using alkaline phosphatase substrate until purple coloration appeared (15–30 min). Plates were washed with tap water and plates were dried in the dark. ELISpot Plate reader… was used for imaging and counting.

## Acknowledgements

We thank the staff at the University of York’s Bioscience Technology Facility Genomics and I&C labs, as well as the Data Science Hub, for their technical support. We also appreciate the staff at the University of York’s and LSHTM Biological Services Facilities for their technical support and animal husbandry. We thank Hernán Carrasco for sharing parasite strains. We thank Shaye Hagler and Jennifer Brooks from Canopy Biosciences (Bruker) for technical support with CellScape ChipCytometry. D.P.-M. extends special thanks to **Jean Langhorne** and **Anne O’Garra** for inspiring discussions during the early conceptualization of this work. This work was funded by a Hull York Medical School Immunology Fellowship, a Royal Society University Research Fellowship (URF\R1\221561), a British Society for Immunology Career Enhancing grant and a European RNAscope™ Genomics Grant (Bio-Techne) awarded to D.P.-M., and an MRC New Investigator Research Grant (MR/R021430/1) awarded to M.D.L. SD, HA, NSD and PMK were supported by a Wellcome Investigator Award (#WT224290 to P.M.K.). A.D. was supported by a Hull York Medical School PGR Scholarship. Figures 1a and 6d-f in this article were created using Biorender.com.

## Author Contributions

D.P.-M. conceptualized, supervised, and coordinated the study. D.P.-M. secured funding with support from P.M.K., J.M.K., and S.A.T. A.D. and D.P.-M. designed the experiments with support from P.M.K., E.A.G., O.A.B., and S.A.T. A.D. and D.P.-M. led data integration and interpretation with support from P.M.K. and E.A.G. A.D. and D.P.-M. performed Visium spatial transcriptomics, single-cell RNA sequencing, ChipCytometry, and immunofluorescence. A.D. carried out computational data analyses with guidance from S.D. on computational pipelines. A.D., A.F.F., A.A.K., K.V.B., K.P., M.D.L., and D.P.-M. conducted mouse infection experiments and tissue collection. S.A., N.S.D., H.A., and K.P. performed RNAscope, ChipCytometry, and tissue imaging assays. K.V.B. contributed to early setup of spatial and single-cell workflows. O.A.B. and S.A.T. provided conceptual advice on experimental design. M.C.T., J.M.K., and M.D.L. provided reporter parasite strains and institutional support. E.A.G., P.M.K., J.M.K., and M.D.L. supervised co-authors and assisted with data interpretation. J.M.K., M.D.L., and P.M.K. contributed funding and reagents. A.D. and D.P.-M. prepared the manuscript with reviewing and editing input from all authors.

---

**Supplementary Figure 1.**
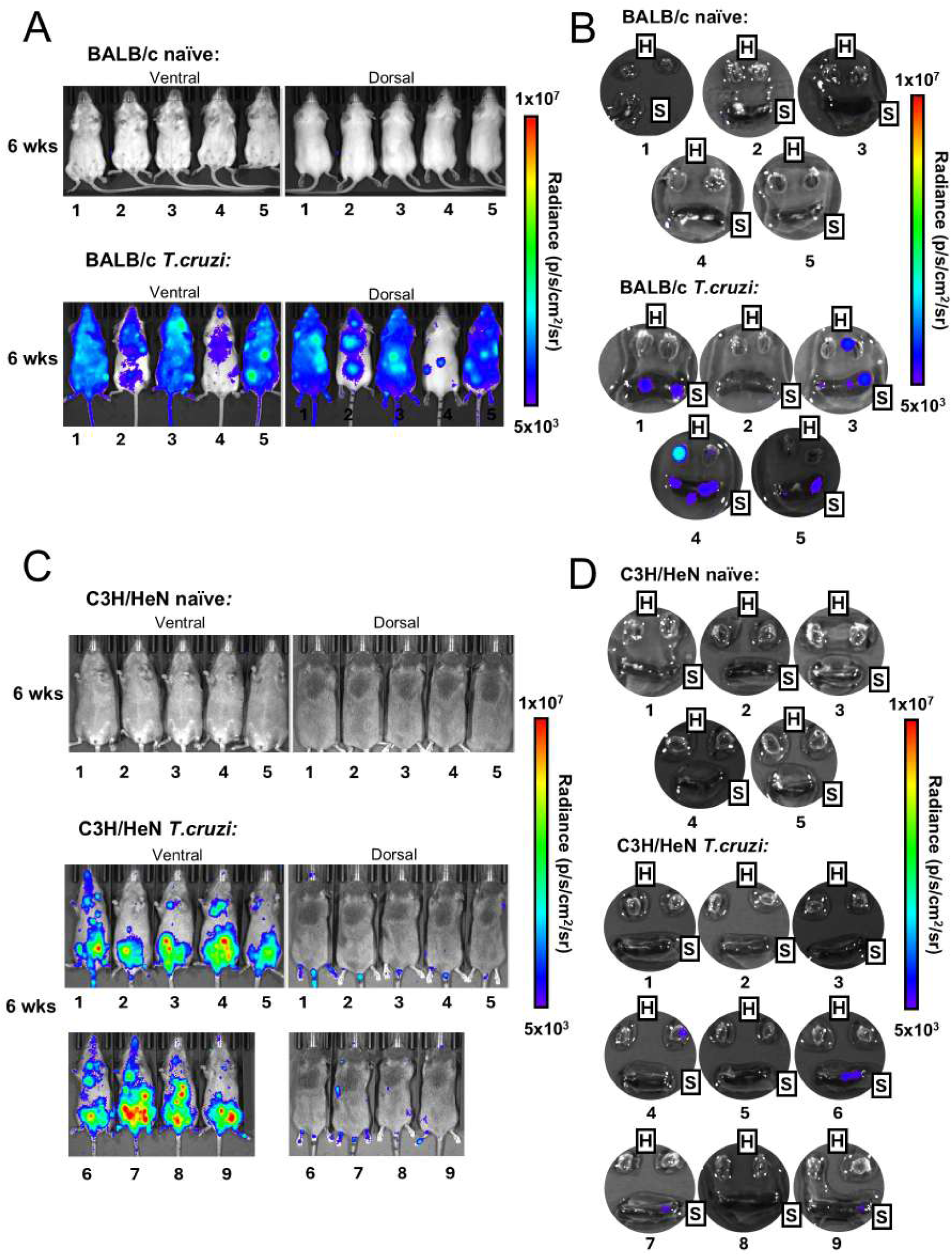
IVIS imaging of parasite burden. *In-vivo* whole-body ventral and dorsal IVIS imaging in BALB/c **A)** and C3H/HeN **C)** mice at 6 weeks p.i. and naïve age-matched controls. *Ex-vivo* organ imaging (heart (H) cross-sectioned into two halves, spleen (S)) in BALB/c **B)** and C3H/HeN **D)** mice at 6 weeks p.i. and naïve age-matched controls.

**Supplementary Figure 2.**
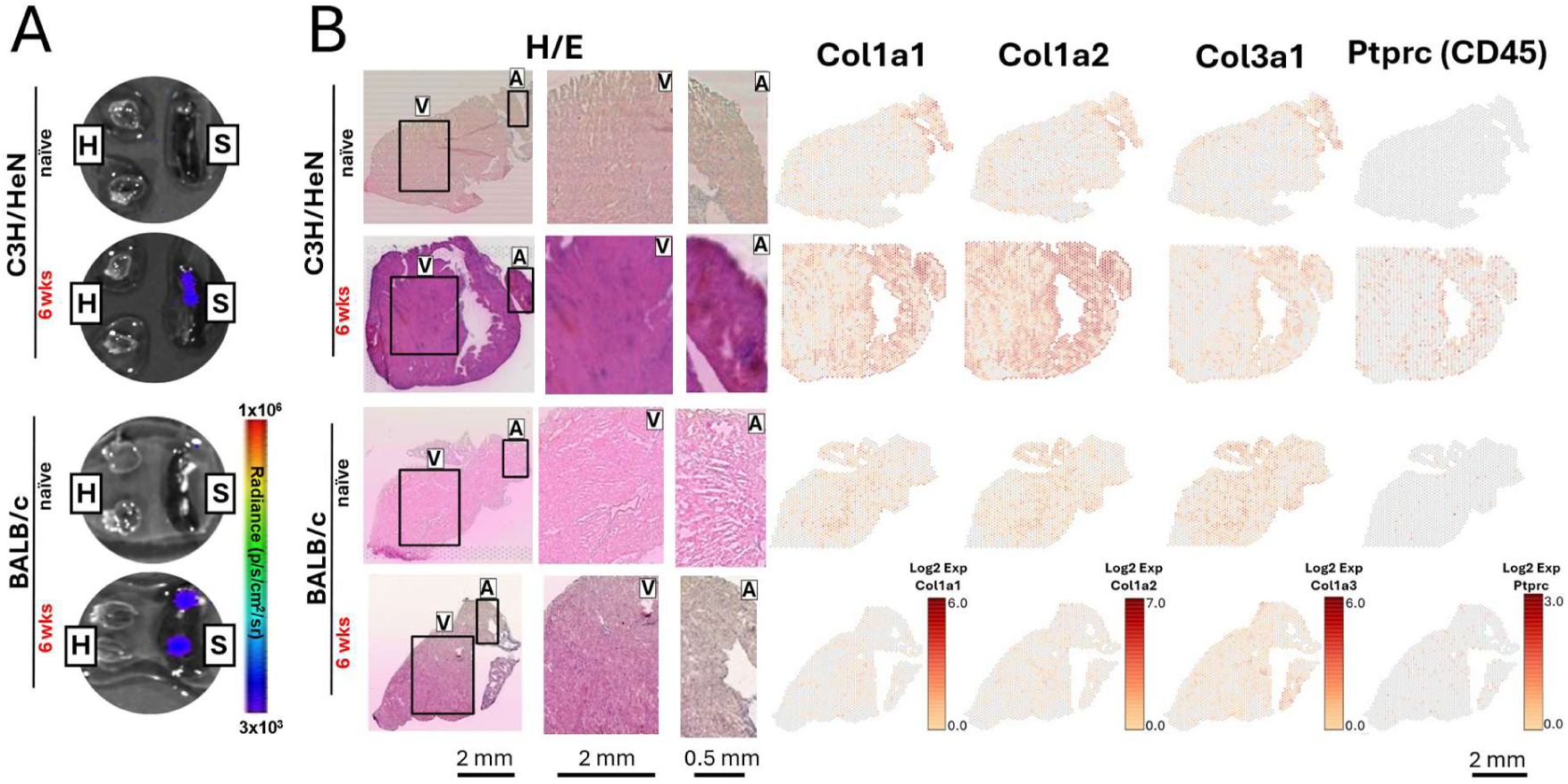
Collagen deposition and immune infiltration in chronic infection. **A)** Representative IVIS images showing parasite burden in harvested organs (heart, spleen) at 6 weeks p.i. **B)** Representative H&E staining displaying the full cardiac cross-section with zoomed views of the ventricle (V) and atrium (A), alongside spatial transcriptomic maps showing log₂ expression of Col1a1/Col1a2 (collagen I), Col3a1 (collagen III), and Ptprc (CD45) across cardiac tissue in each condition.

**Supplementary Figure 3.**
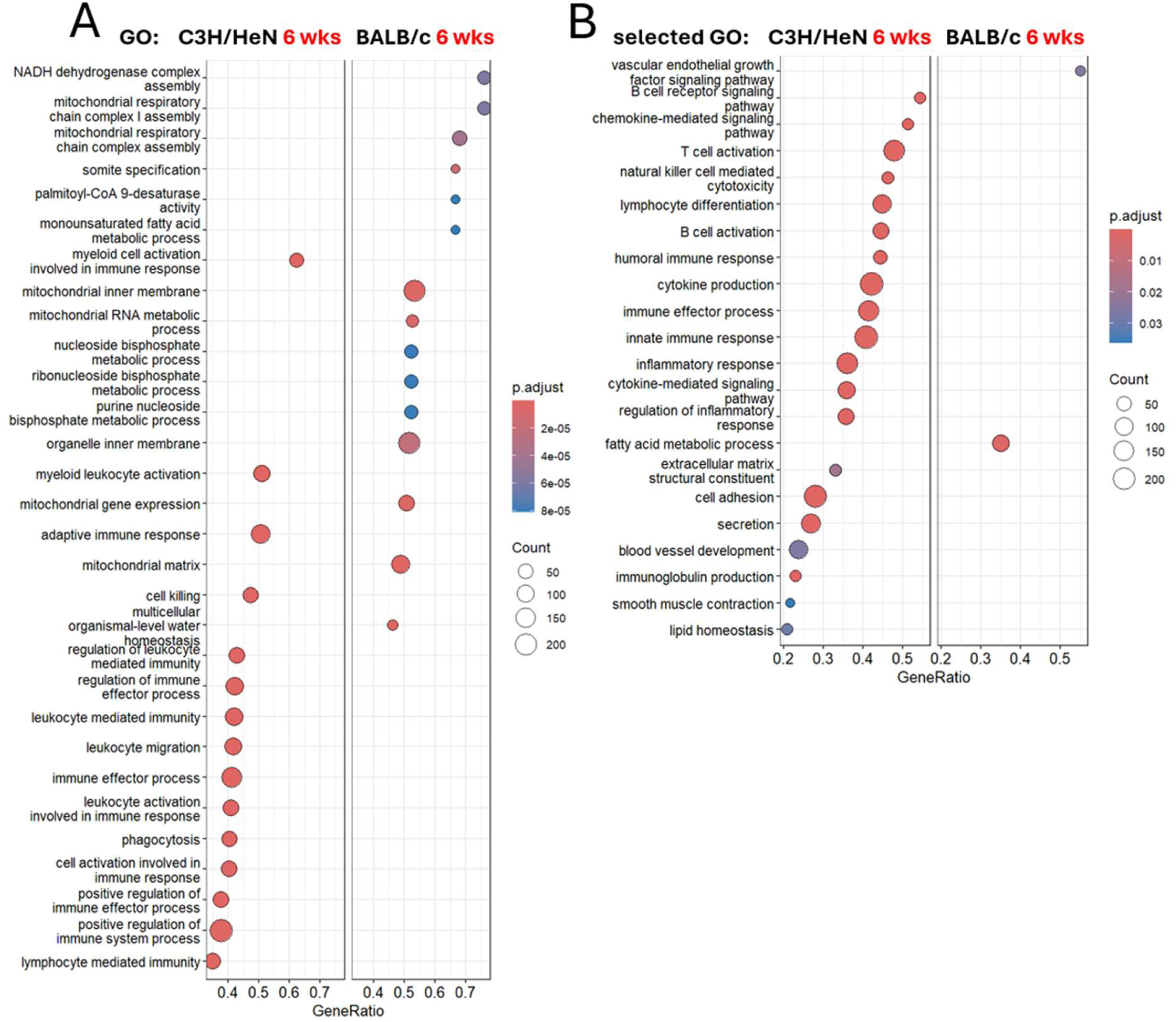
GO analysis of spatial transcriptomic data. **A)** Top 30 biological processes (ranked by adjusted p-value) identified through GO analysis comparing C3H/HeN and BALB/c mice at 6 weeks p.i. **B)** Selected immune-related GO processes enriched between strains at 6 weeks p.i.

**Supplementary Figure 4.**
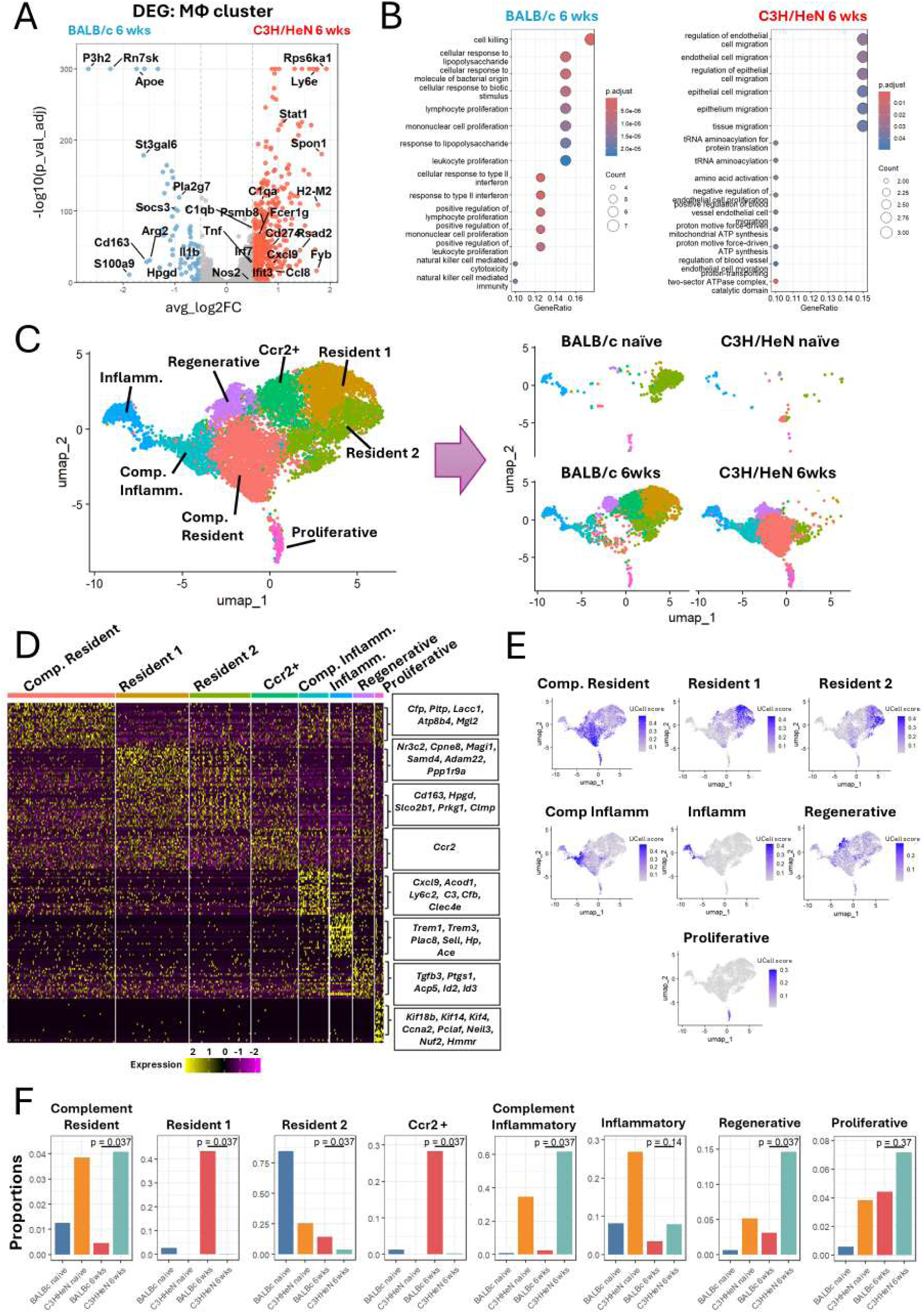
Transcriptional profiling of cardiac macrophages during chronic infection. **A)** Volcano plot showing differential gene expression in macrophages between C3H/HeN and BALB/c mice at 6 weeks p.i. **B)** Top 15 enriched biological processes (ranked by adjusted *P* value) identified by Gene Ontology analysis for C3H/HeN 6 weeks vs naïve and BALB/c 6 weeks vs naïve comparisons. **C)** UMAP of sub-clustered cardiac macrophages, highlighting the major macrophage states identified by scRNA-seq. **D)** Heatmap showing the top 25 differentially expressed genes for each macrophage sub-cluster. Highlighted genes were used for UCell scoring. **E)** Feature plots showing UCell scores for the identified macrophage states across the macrophage UMAP. **F)** Mean proportions of macrophage subpopulations relative to total macrophages. *P* values indicate two-sided Wilcoxon rank-sum tests comparing BALB/c 6 weeks and C3H/HeN 6 weeks samples.

**Supplementary Figure 5.**
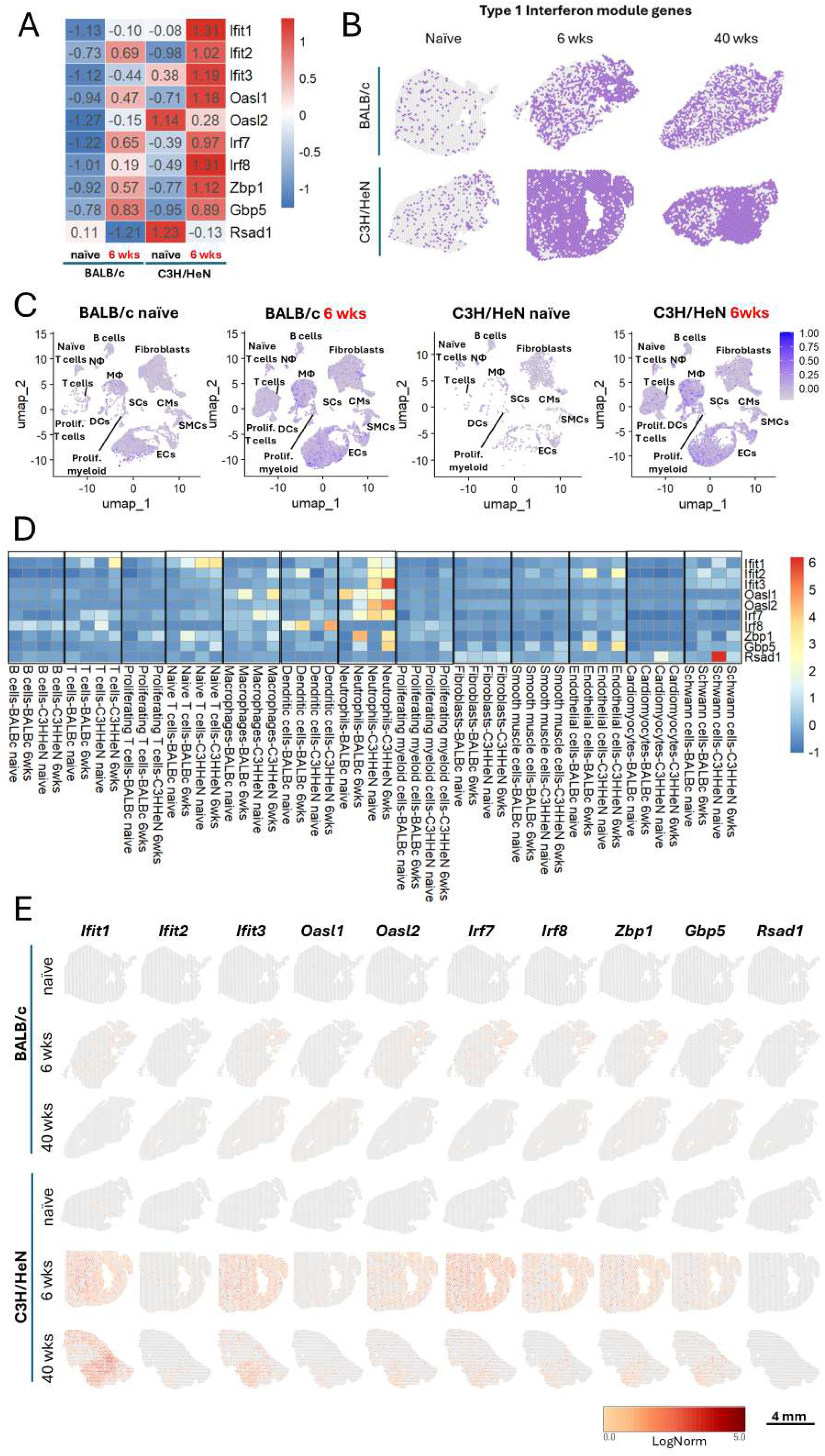
Type I interferon gene expression in the heart. **A)** Heatmap showing the expression (z-scores) of Type I interferon genes across conditions in the scRNA-seq heart dataset. **B)** Representative spatial transcriptomic plots showing heart cross-sections with spots positive for either of the Type I interferon module genes (purple). **C)** UMAP plots showing the expression of Type I interferon module genes across the identified cell clusters in the scRNA-seq heart dataset. **D)** Heatmap showing the expression (z-scores) of Type I interferon genes across cell types and conditions in the scRNA-seq heart dataset. **E)** Representative spatial transcriptomic plots showing the log-normalized expression of individual Type I interferon genes in heart cross-sections from naïve, 6-week, and 40-week BALB/c and C3H/HeN mice.

**Supplementary Figure 6.**
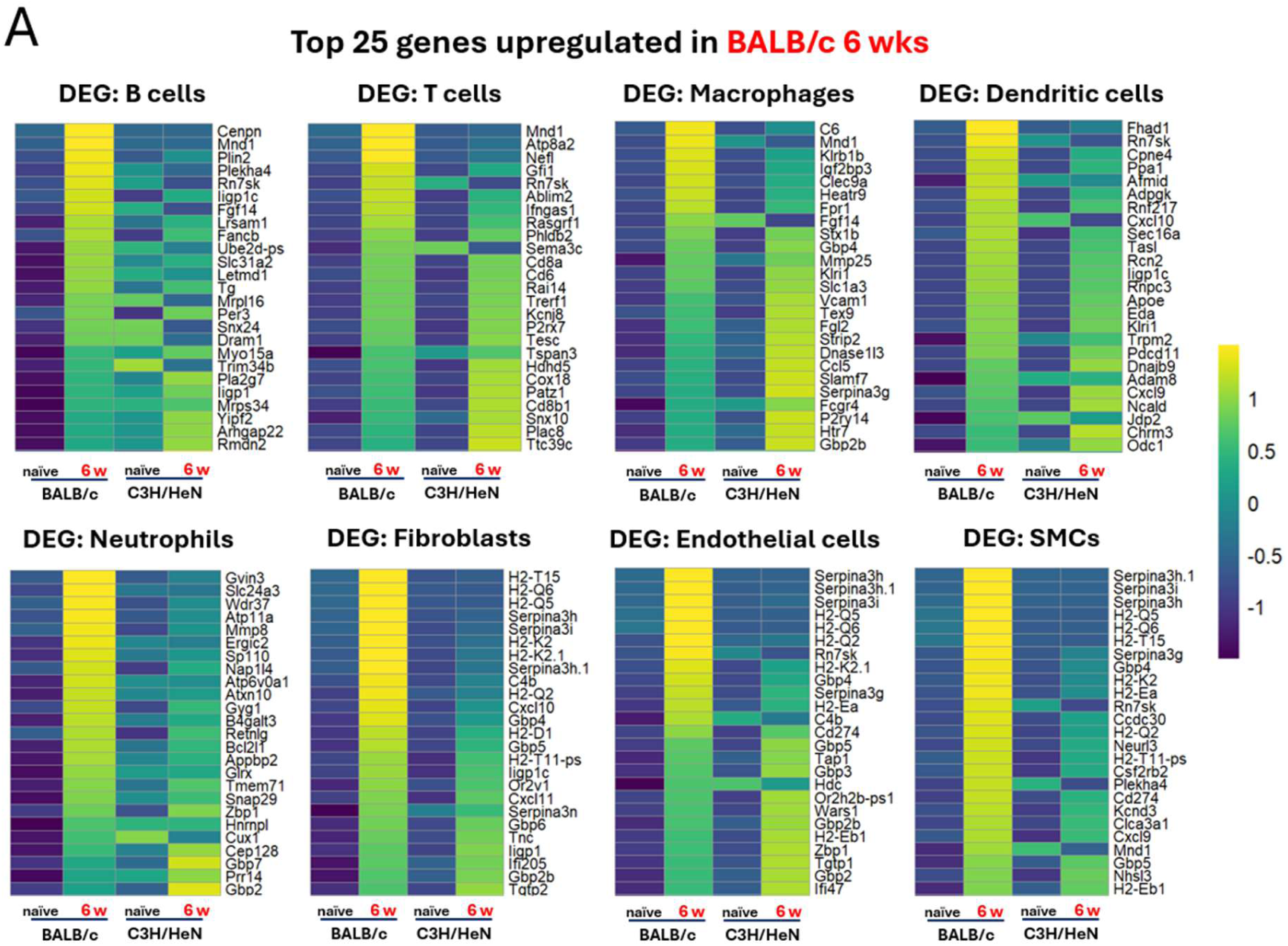
BALB/c differential expression heatmaps. **A)** Heatmaps showing the top 25 highest z-scores in selected cell types for BALB/c at 6 weeks p.i. compared with other conditions. Genes were selected from the top 50 upregulated genes in BALB/c 6 weeks relative to BALB/c naïve.

**Supplementary Figure 7.**
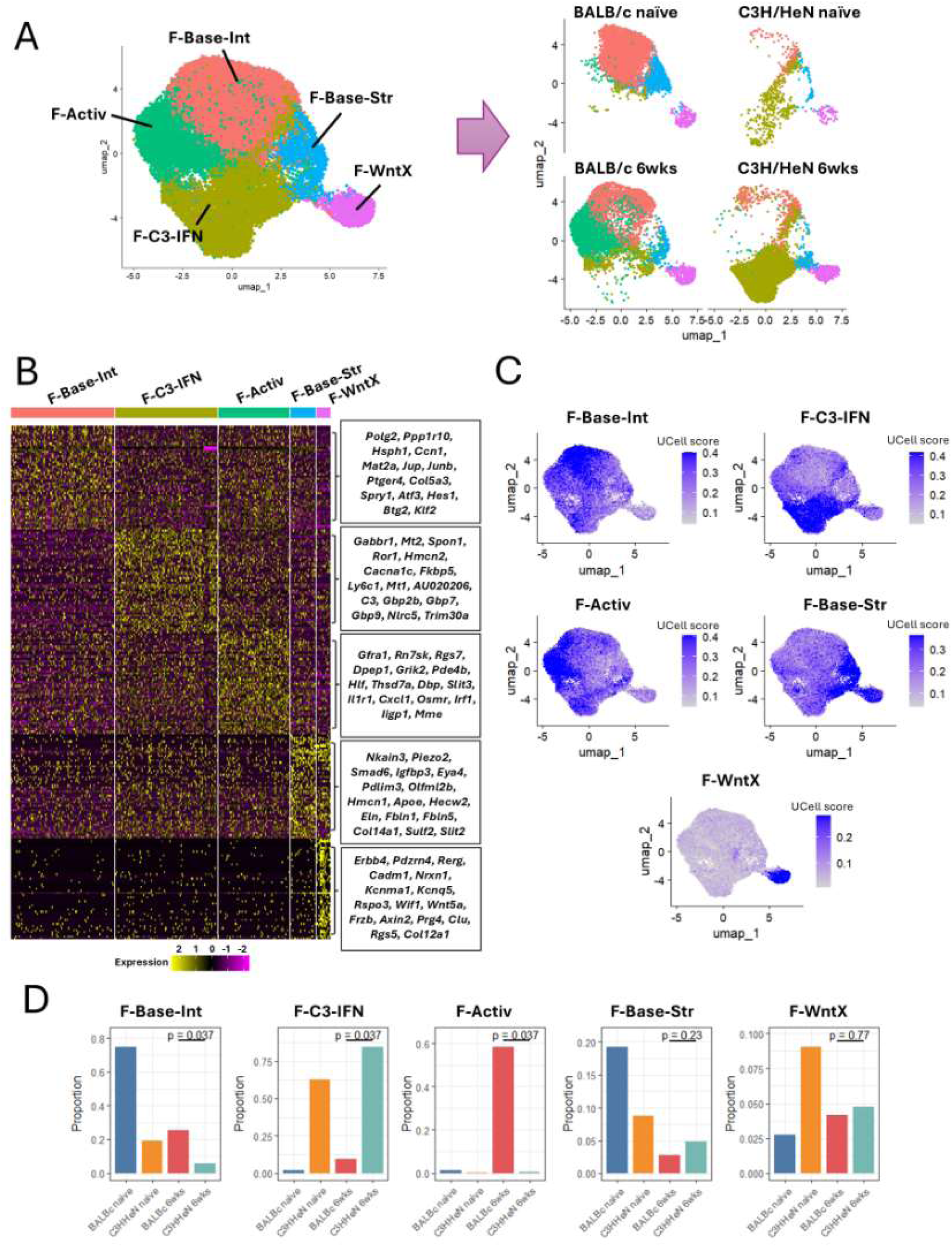
Transcriptional profiling of fibroblasts during chronic infection. **A)** UMAP of sub-clustered cardiac fibroblasts, highlighting the major fibroblast populations identified by scRNA-seq. **B)** Heatmap showing the top 50 differentially expressed genes for each fibroblast sub-cluster. Highlighted genes were used for UCell scoring. **C)** Feature plots showing UCell scores for the identified fibroblast populations across the fibroblast UMAP. **D)** Mean proportions of fibroblast populations relative to total fibroblasts. *P* values indicate two-sided Wilcoxon rank-sum tests comparing BALB/c 6 weeks and C3H/HeN 6 weeks samples.

**Supplementary Figure 8.**
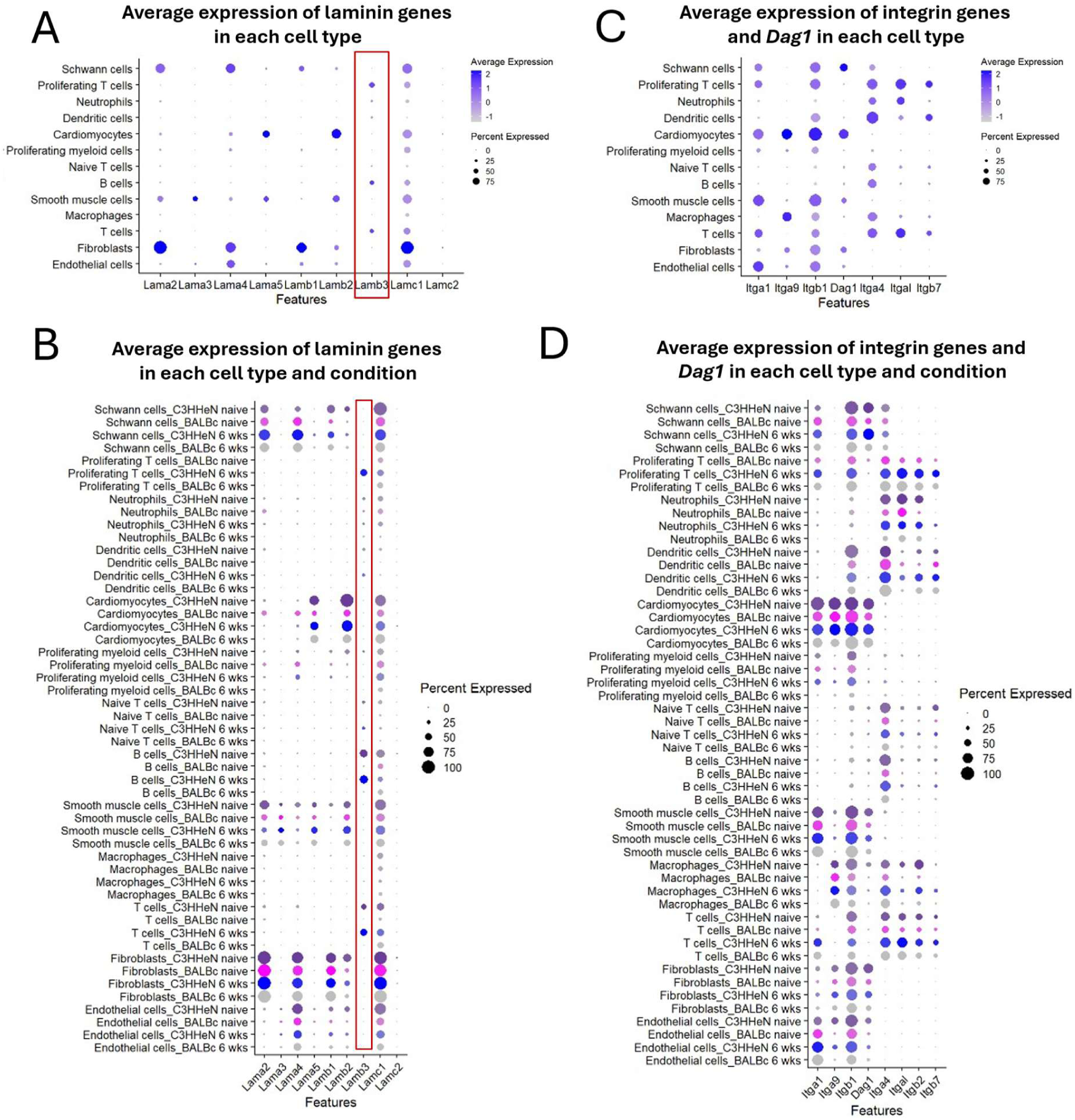
Laminin, integrin, and *Dag1* expression in the scRNA-seq heart dataset. **A)** Average expression of laminin genes in each cell type. **B)** Average expression of laminin genes in each cell type and condition. **C)** Average expression of integrin genes and *Dag1* in each cell type. **D)** Average expression of integrin genes and *Dag1* in each cell type and condition.

**Supplementary Figure 9.**
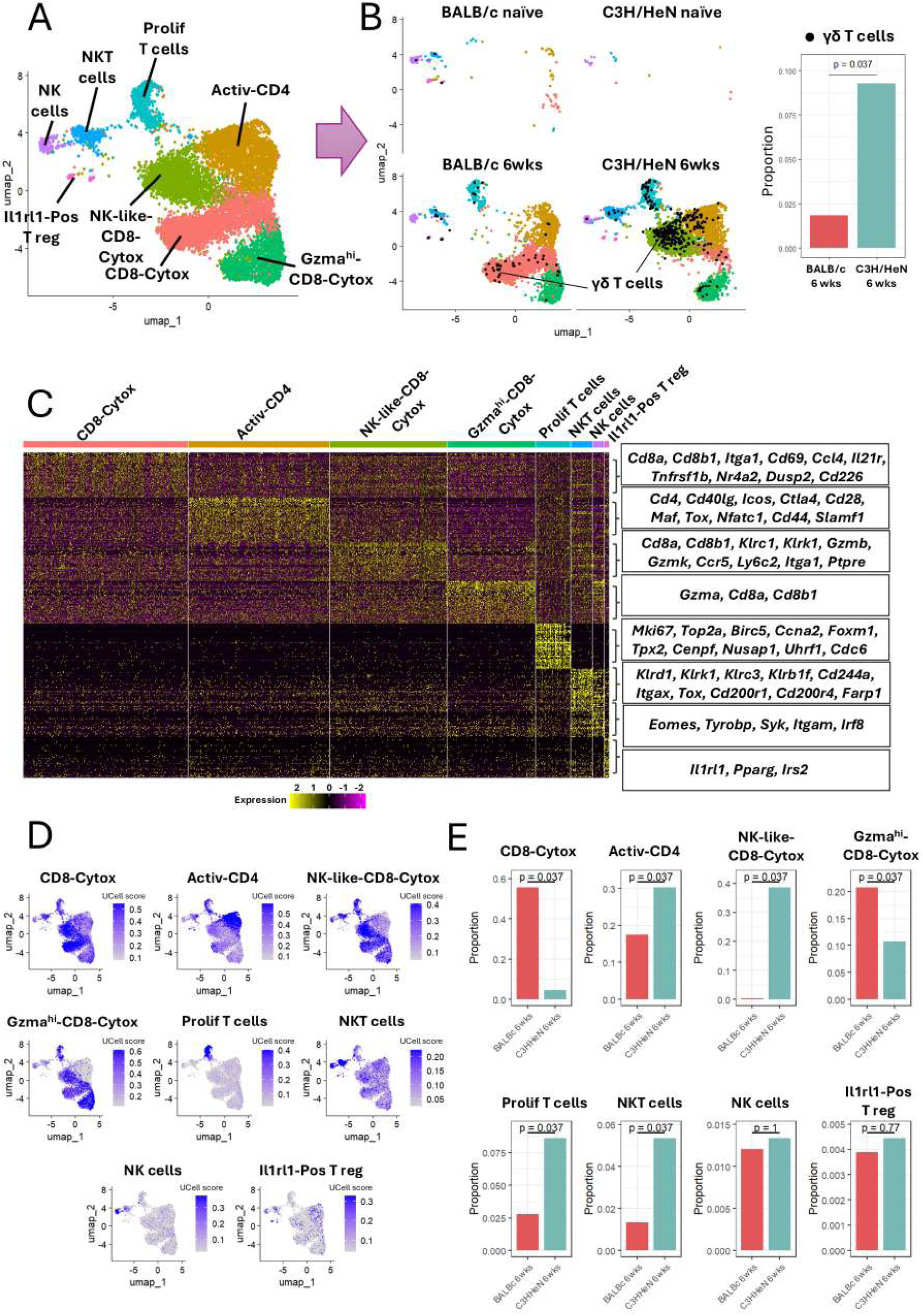
Transcriptional profiling of T cells during chronic infection. **A)** UMAP of sub-clustered cardiac T cells, highlighting the major T cell populations identified by scRNA-seq. **B)** UMAPs of sub-clustered cardiac T cells split by condition, with identified γδ T cells overlaid in black. Bar plot showing the mean proportion of γδ T cells relative to total T cells. *P* values indicate two-sided Wilcoxon rank-sum tests comparing BALB/c and C3H/HeN mice at 6 weeks. **C)** Heatmap showing the top 50 differentially expressed genes for each T cell sub-cluster. Highlighted genes were used for UCell scoring. **D)** Feature plots showing UCell scores for the identified T cell populations across the T cell UMAP. **E)** Mean proportions of T cell populations relative to total T cells. *P* values indicate two-sided Wilcoxon rank-sum tests comparing BALB/c 6 weeks and C3H/HeN 6 weeks samples.

